# Cell-type-specific cholinergic control of granular retrosplenial cortex with implications for angular velocity coding across brain states

**DOI:** 10.1101/2024.06.04.597341

**Authors:** Izabela Jedrasiak-Cape, Chloe Rybicki-Kler, Isla Brooks, Megha Ghosh, Ellen K.W. Brennan, Sameer Kailasa, Tyler G. Ekins, Alan Rupp, Omar J. Ahmed

**Affiliations:** Dept. of Psychology, University of Michigan, Ann Arbor, MI 48109; Neuroscience Graduate Program, University of Michigan, Ann Arbor, MI 48109; Kresge Hearing Research Institute, University of Michigan, Ann Arbor, MI 48109; Dept. of Biomedical Engineering, University of Michigan, Ann Arbor, MI 48109; Dept. of Mathematics, University of Michigan, Ann Arbor, MI 48109; Dept. of Internal Medicine, University of Michigan, Ann Arbor, MI 48109

## Abstract

Cholinergic receptor activation enables the persistent firing of cortical pyramidal neurons, providing a key cellular basis for theories of spatial navigation involving working memory, path integration, and head direction encoding. The granular retrosplenial cortex (RSG) is important for spatially-guided behaviors, but how acetylcholine impacts RSG neurons is unknown. Here, we show that a transcriptomically, morphologically, and biophysically distinct RSG cell-type – the low-rheobase (LR) neuron – has a very distinct expression profile of cholinergic muscarinic receptors compared to all other neighboring excitatory neuronal subtypes. LR neurons do not fire persistently in response to cholinergic agonists, in stark contrast to all other principal neuronal subtypes examined within the RSG and across midline cortex. This lack of persistence allows LR neuron models to rapidly compute angular head velocity (AHV), independent of cholinergic changes seen during navigation. Thus, LR neurons can consistently compute AHV across brain states, highlighting the specialized RSG neural codes supporting navigation.

## INTRODUCTION

Acetylcholine (ACh) influences neuronal excitability, circuit dynamics, synaptic transmission, and plasticity across the cortex (Conner et al., 2003, 2010; Gil et al., 1997; Hasselmo, 1995, 1999; Hasselmo & Giocomo, 2006; Jiang et al., 2016; Ljubojevic et al, 2018; Mitsushima et al., 2013; Muñoz and Rudy, 2014; Nakajima et al., 1986; Picciotto et al., 2012; Rasmusson, 2000; Sarter et al., 2009; Sarter and Lustig, 2020; Verhoog et al., 2016). Cholinergic transmission is important for attention and goal-directed behaviors (Arnold et al., 2002; Hasselmo, 2006; Luchicchi et al., 2014; Sarter et al., 2014, 2005), including spatial navigation (Hamlin et al., 2013; Solari and Hangya, 2018; Stancampiano et al., 1999; Whishaw, 1985; Winkler et al., 1995; Zannone et al., 2018). To aid in successful navigation, ACh levels increase throughout spatially-relevant regions (Anzalone et al., 2009; Giovannini et al., 2001; Sarter et al., 2005) to induce cortical synaptic plasticity (Kilgard and Merzenich, 1998; Pinto et al., 2013; Rasmusson, 2000; Zannone et al., 2018), increase sensitivity to behaviorally relevant cues (Parikh et al., 2007; Pinto et al., 2013; Sarter et al., 2005), and facilitate sensorimotor gating (Jin et al., 2019; Luntz-Leybman et al., 1992; Sarter et al., 2005) . Blocking cholinergic transmission in the hippocampus and cortex results in significantly impaired performance on spatial tasks (Berger-Sweeney et al., 1994; Hamlin et al., 2013), highlighting the key role cholinergic inputs play in navigation.

At the cellular level, ACh induces persistent firing of principal neurons in spatially relevant brain regions, including the entorhinal cortex (Jochems et al., 2013; Tahvildari et al., 2007; Yoshida et al., 2008), CA1 (Knauer et al., 2013; Yamada-Hanff and Bean, 2013), CA3 (Bianchi and Wong, 1994; Jochems and Yoshida, 2013) dorsal subiculum (Kawasaki et al., 1999), postsubiculum (Yoshida and Hasselmo, 2009), motor cortex (Rahman and Berger, 2011), anterior cingulate cortex (ACC; Ratté et al., 2018; Zhang and Seguela, 2010), primary visual and somatosensory cortices (Rahman and Berger, 2011), and prefrontal cortex (Dembrow et al., 2010; Gulledge et al., 2009; Gulledge, 2024; Thuault et al., 2013). In many cortical areas, cholinergic induced persistent activity is seen across layers, in both superficial (Navaroli et al., 2012; Ratté et al., 2018; Tahvildari et al., 2007; Yoshida et al., 2008; Zhang and Seguela, 2010) and deep (Egorov et al., 2002; Fu et al., 2019; Gulledge et al., 2009; Gulledge, 2024; Rahman and Berger, 2011; Thuault et al., 2013) pyramidal neurons, consistent with cholinergic projections spanning cortical layers (Eckenstein et al., 1988; Levey, 1993; Vogt, 1984).

The basal forebrain (BF) is one of the main hubs of cholinergic neurons in the brain (Mesulam et al., 1983) and sends widespread projections to the cortex to influence cortical function (Hsieh et al., 2000; Kawaguchi, 1997; Sarter et al., 2005; Sarter and Lustig, 2020; Verhoog et al., 2016; Yuan et al., 2019). One key target of these projections is the retrosplenial cortex (RSC), a navigationally important region which receives cholinergic inputs in both superficial and deep layers from multiple parts of the BF (van Groen and Wyss, 2003, 1992, 1990). The granular region of the RSC (RSG) also receives converging input from navigationally-important regions including the dorsal subiculum (Brennan et al., 2021; Opalka and Wang, 2020; Wyss and van Groen, 1992; Yamawaki et al., 2019a), anterior thalamus (Brennan et al., 2021; Lomi et al., 2021; van der Goes et al., 2024; Yamawaki et al., 2019b), as well as inhibitory inputs from the CA1 of the hippocampus (Yamawaki et al., 2019b). RSG also shows a larger increase in concentration of ACh during navigation compared to both the dysgranular retrosplenial cortex (RSD) and the hippocampus (Anzalone et al., 2009). However, despite the convergence of multiple cholinergic inputs onto RSG cells and the dramatic increase in ACh release within RSG during navigation, how cholinergic agonists impact RSG spatial computations remains unknown.

The RSG contains multiple principal neuronal subtypes, including a hyperexcitable small pyramidal neuron called the low rheobase cell (LR; Brennan et al., 2020). LR cells constitute the most numerous principal neuronal subtype in the RSG, but are not found in other cortical regions (Brennan et al., 2020., Brennan et al., 2021, Kurotani et al., 2013., Robles et al., 2020, Sullivan et al., 2023, Wyss et al., 1990, Yousuf et al., 2020). Here, using a combination of single cell transcriptomics, whole-cell physiology, pharmacology, and biophysical modeling we uncover how cholinergic agonists control distinct principal pyramidal cell types within the RSG, RSD and frontal cortices. We find that LR cells represent a remarkably distinct principal neuronal subtype compared to all other examined neurons, exemplified by radically different responses to cholinergic agonists. Computational modeling shows that this allows LR cells to rapidly compute the derivative of head direction – angular head velocity – independent of cholinergic tone. Thus, the principal neurons that selectively receive the strongest directional and spatial inputs from anterior thalamus and dorsal subiculum — LR neurons (Brennan et al., 2021) — are also the ones that never fire persistently in response to cholinergic stimulation. This highlights a non-canonical coding schema in RSG that can support rapid, algorithmically consistent computations despite ongoing changes in cholinergic state between quiet wakefulness and active navigation.

## RESULTS

### Low Rheobase cells have a distinct cholinergic receptor transcriptomic signature

We performed single-nuclei RNA sequencing (snRNA-seq) on samples of microdissected mouse RSG using the 10x Chromium technique. Unsupervised clustering of the ∼8,500 cells that passed quality control metrics (see Methods) yielded several distinct clusters. To identify the cell types that these clusters correspond to, we utilized a published 10x RNA-seq dataset (Yao et al. 2021) as a reference dataset for canonical-correlation analysis (CCA). Each cell was assigned a cluster label from the more than 300 identified in the reference dataset. From these labels, we were able to define putative cell types for the vast majority of our clusters. The reference dataset additionally proposes a cortical layer range for each glutamatergic cell type, allowing us to perform cortical layer-specific transcriptomic analyses.

More than one-third of all identified neuronal cells in our dataset fall into a glutamatergic, putatively L2/3 cluster denoted “L2/3 Cxcl14” due to high expression levels of the gene *Cxcl14*. Within L2/3, 84% of excitatory neurons fall within this cluster (Figure 1B, left), which matches our earlier physiological data on neuronal subtype density in superficial layers of RSG. Specifically, we have previously shown that 81% of physiologically recorded L2/3 RSG pyramidal neurons are LR neurons (Figure 1B and Brennan et al., 2020). We used Fisher’s exact test to assess the validity of our assumption that L2/3 physiologically defined LR and RS cells correspond to the transcriptomic L2/3 Cxcl14 and L2/3 Calb1 clusters, respectively. Under this assumption, we cannot reject the null hypothesis that the transcriptomic and physiological data were drawn from the same distribution (p=0.48). However, if we assume that the transcriptomic clusters are flipped, we can reject the null hypothesis (p = 8.0e-49). This strongly suggested that the L2/3 Cxcl14 cluster represents LR cells in RSG. To further confirm this, we consulted in-situ hybridization (ISH) data for the *Cxcl14* gene from the Allen Brain Atlas database (Lein et al., 2007), which reveals dense labeling of cortical layers 2 and 3 in RSG that mirrors the distribution of LR neurons in RSG (Brennan et al., 2020; Robles et al., 2020). Thus, the localization of *Cxcl14*, the size of this cluster compared to the total number of all L2/3 excitatory cells in our dataset, and previous anatomical and physiological observations together point to physiological LR neurons corresponding to our L2/3 Cxcl14 cluster. This, in conjunction with other studies (Brennan et al., 2020; Sullivan et al., 2023; Yao et al., 2021), argues that L2/3 Cxcl14 cells are physiological LR cells, while L2/3 Calb1 cells are physiological RS cells.

**Figure 1.**
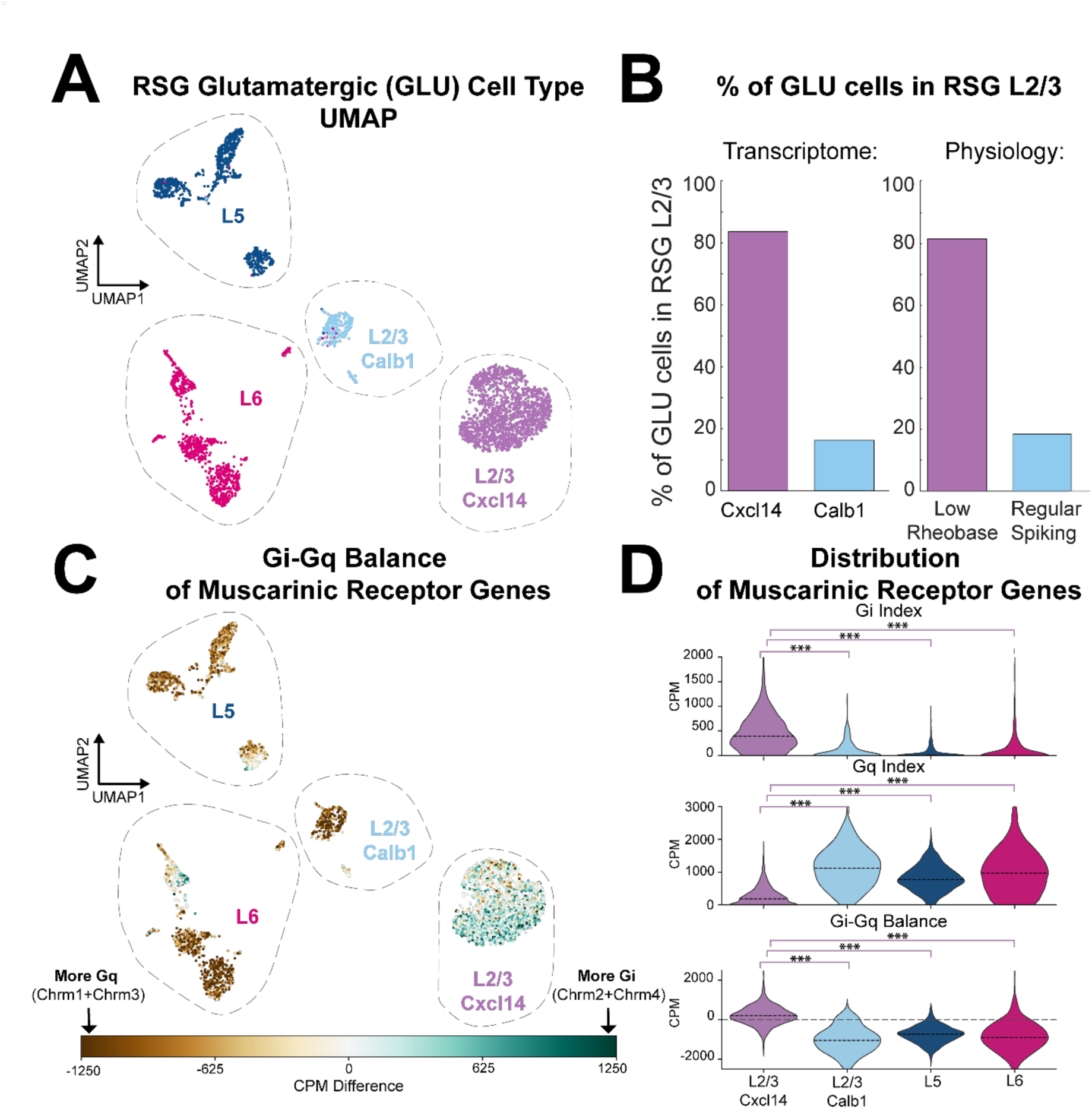
Retrosplenial low rheobase (LR) cells have a dramatically different cholinergic receptor (Chrm) expression pattern compared to other glutamatergic cell types in RSG. ***A.*** UMAP of glutamatergic single-nuclei RNA sequenced RSG cells grouped by layer. Layer 2/3 neurons were further subdivided to reveal two distinct clusters defined by the expression of *Cxcl14* and *Calb1* genes, highlighting two separate excitatory cell populations in the superficial layers of RSG. ***B.*** Relative percentage of the two distinct glutamatergic cell populations in RSG layer 2/3 grouped by transcriptome (based on clustering from A) and by physiology (based on previously published experimental data: Brennan et al., 2020). Note a near-identical proportion of transcriptomically identified Cxcl14 RSG neurons and physiologically identified LR neurons, mapping the transcriptomic L2/3 Cxcl14 cluster onto the physiological LR neurons. ***C.*** UMAP from (A), colored by difference in counts per million (CPM) between functionally distinct G protein coupled muscarinic acetylcholine receptor genes (*Chrm1-4*). Gi represents the sum of CPM of *Chrm2* + *Chrm4*, which are coupled to Gαi receptor subunit and show largely hyperpolarizing modulation. Gq represents the sum of CPM of *Chrm1* + *Chrm3*, which are coupled to Gαq receptor subunit and show depolarizing modulation. The L2/3 Cxcl14 cluster (representing physiological LR cells) is distinctly skewed towards Gαi-coupled receptor expression. ***D.*** Violin plots of Chrm gene expression in each analysis category. Gi and Gq definitions are the same as in **C**. Gi-Gq Balance quantifies the difference shown in **C**. Expression is significantly different between L2/3 Cxcl14 (LR cells) and all other clusters (*** p < .001). This suggests that LR neurons should show a distinct response to cholinergic activation, relative to cells in other clusters.

We next examined the expression pattern for muscarinic cholinergic receptor (mAChR) genes across transcriptomic clusters. We focused on four well established classes of this receptor expressed in the brain, M1-M4 (Caulfield and Birdsall, 1998), corresponding to the *Chrm1-Chrm4* genes. All muscarinic receptors are members of the G protein coupled receptor family but M1 and M3 couple to Gαq subunit, while M2 and M4 couple to Gαi subunit. This distinction correlates with downstream cellular effects. While the neuromodulatory action of acetylcholine via muscarinic receptors in the cortex is complex (Muñoz and Rudy, 2014), Gαq-coupled M1 and M3 generally increase the excitability of cells (Caulfield, 1993; Caulfield and Birdsall, 1998). On the other hand, Gαi-coupled M2 and M4 preferentially lead to hyperpolarization of their targets, and these receptors are often expressed on presynaptic axon terminals where they can help to suppress neurotransmitter release (Egan and North, 1986; Caulfield and Birdsall, 1998). Thus, we summed M1 and M3 gene expression (*Chrm1 + Chrm3*) under the “Gq Index” label and summed the M2 and M4 gene expression (*Chrm2 + Chrm4*) under the “Gi Index” label (see methods), although we noted that *Chrm4* was very weakly expressed throughout this dataset. Examining the expression profiles in the physiological LR cluster (transcriptomic L2/3 Cxcl14 cluster) highlighted stark differences compared to all other glutamatergic clusters in RSG. Specifically, LR cells express far more of the canonically inhibitory M2 and M4 mAChR genes (Gi Index, effect size: L2/3 Cxcl14 vs L2/3 Calb1 Cohen’s *d = 0.97*; L2/3 Cxcl14 vs L5 Cohen’s *d =* 1.14; L2/3 Cxcl14 vs L6 Cohen’s *d =* 0.94, all p < 0.001), and far less of the canonically excitatory M1 and M3 mAChRs (Gq Index, effect size: L2/3 Cxcl14 vs L2/3 Calb1 Cohen’s *d = 1.85*; L2/3 Cxcl14 vs L5 Cohen’s *d =* 1.34; L2/3 Cxcl14 vs L6 Cohen’s *d =* 1.26, all p < 0.001) than other glutamatergic cell types. An analysis of the difference between those two indices (Gi – Gq Balance) yields similarly significant results (effect size: L2/3 Cxcl14 vs L2/3 Calb1 Cohen’s *d = 1.82*; L2/3 Cxcl14 vs L5 Cohen’s *d =* 1.49; L2/3 Cxcl14 vs L6 Cohen’s *d =* 1.36, all p < 0.001) (Figures 1A, 1C, and 1D).

These results led us to hypothesize that direct cholinergic control of excitatory cells in RSG may be cell-type-specific, with physiological LR cells potentially showing very different responses from other principal excitatory neuronal subtypes. To test this hypothesis, we next examined the physiological modulation of individual RSG cells by cholinergic agonists.

### Cholinergic agonists induce persistent activity in RS, but not LR, neurons in the granular retrosplenial cortex

Using whole-cell patch clamp, we recorded from four previously recognized physiological pyramidal cell populations in layers 2-5 of the granular retrosplenial cortex (RSG; Brennan et al., 2020, 2021; Yousuf et al., 2020). As in previous work, (Brennan et al., 2020, 2021) we identified these cell types by their intrinsic, spiking, and anatomical properties: L2/3 LR neurons, L3 RS neurons, L5 RS neurons, and L5 intrinsically bursting (L5 IB) neurons. The superficial LR and RS neurons correspond to the L2/3 Cxcl14 and L2/3 Calb1 transcriptomic clusters, respectively. Physiologically, LR cells differed from neighboring L3 RS neurons based on their significantly lower rheobase, higher input resistance, and lower spike frequency adaptation (Figure 2A, Supplemental Table 1, Brennan et al., 2020). LR cells also had clear morphological differences from all recorded RS cells, with LRs having a smaller soma size and sparse dendritic branching pattern (Figure 2A; Brennan et al., 2020, 2021). L5 IB cells were distinguished from L5 RS neurons by prominent bursts that occur at near-threshold current injections (Figure 2A; Supplemental Table 1).

**Figure 2.**
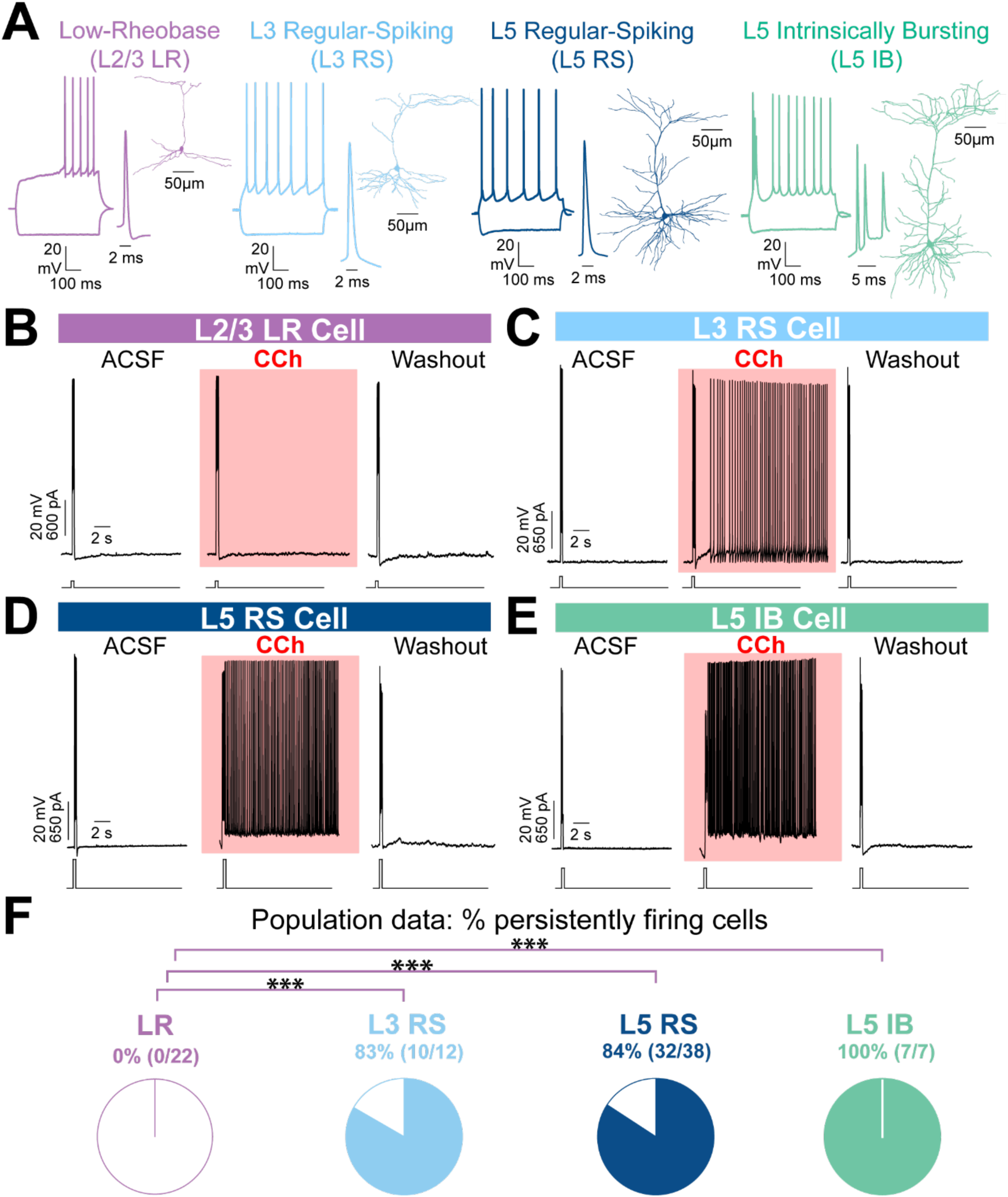
Persistent firing is a cell type-specific activity present in both superficial and deep layers of the RSG. LR cells constitute the only glutamatergic cell-type that shows a complete absence of persistent firing. ***A.*** Representative reconstructions and firing properties of LR (purple; left), L3 RS (light blue; middle left), L5 RS (navy; middle right), and L5 IB (teal; right) neurons. ***B.*** Responses from a representative LR neuron (shown in ***A***). Left: Baseline trace of LR activity in normal ACSF. Middle: Trace showing lack of persistent firing after exposure to CCh. Right: Trace of activity after washout of CCh with normal ACSF. The short-lasting current step to trigger the initial firing is shown under each voltage trace. ***C.*** Responses from a representative L3 RS neuron (shown in ***A***). Left: Baseline trace of L3 RS activity in normal ACSF. Middle: Trace of persistent firing after exposure to CCh. Right: Trace of activity after washout of CCh with normal ACSF. ***D.*** Same as ***C*** but for a representative L5 RS neuron. ***E.*** Same as ***C*** but for a representative L5 IB neuron. ***F.*** Population analysis of the percentage of neurons within each cell type that exhibited persistent firing. None of the 22 tested LR neurons showed persistence, contrasting sharply with all other principal cell types tested (*** p < 0.001), highlighting cell-type specific cholinergic control of RSG.

To examine how RSG neurons are impacted by cholinergic agonists, we combined whole-cell recordings with bath application of the muscarinic receptor agonist carbachol (CCh). (Figure S1; see Methods). We found that 83-100% of RS and IB neurons fired persistently when exposed to CCh and that responses returned to baseline upon drug washout (Figure 2C-E). There was no significant difference in percentage of cells persistently firing between L3 RS, L5 RS and L5 IB groups (L3 RS v L5 RS: χ^2^(1, n = 50) = 0.005, p = 0.94; L3 RS v L5 IB: χ^2^(1, n = 19) = 1.30, p = 0.25; L5 RS v L5 IB: χ^2^(1, n = 45) = 1.28, p = 0.26; chi-squared test for all). In stark contrast, 0 out of 22 LR neurons fired persistently (Figure 2B&F) making LRs significantly different from all other cell types tested (LR v L3 RS: χ^2^(1, n = 34) = 25.97, p = 3.46e-07; LR v L5 RS: χ^2^(1, n = 60) = 39.7, p = 2.96e-10 ; LR v L5 IB: χ^2^(1, n = 29) = 29.00, p = 7.23e-08; chi-squared test for all). These results indicate that muscarinic receptor activation affects RSG neurons in a cell-type-specific manner, such that LR cells are the only pyramidal cell subtype examined in RSG that do not exhibit persistent activity.

To explore this distinction further, we next investigated another cellular response to cholinergic activation, slow afterdepolarization that precedes persistent firing.

### Slow afterdepolarization is the subthreshold precursor of persistent activity and is absent in LR cells

Neurons that exhibit cholinergic-induced persistent activity first show a gradual build-up of a slow afterdepolarization potential (sADP) that precedes persistent firing (Haj-Dahmane and Andrade, 1998; Navaroli et al., 2012). To more closely examine the persistent activity of these four RSG cell types, we next analyzed sADP amplitudes of all tested neurons. We found that 0/22 LR neurons showed any sADPs, regardless of time exposed to CCh (Figure S2A & F). In contrast, 92% of L3 RS, 95% of L5 RS, and 100% of L5 IB cells exhibited clear sADPs in the sweeps preceding persistent firing (Figure S2B, C, D, F). Even for the few L3 RS, L5 RS, and L5 IB cells that did not fire persistently, a clear trend toward sustained depolarization was observed in the form of sADPs, drawing a clear distinction between LRs and other rare excitatory cells that showed no persistent firing. The lack of sADP in LR neurons was significantly different compared to sADP occurrence in the other 3 cell groups (p < 0.001 for all, chi-squared test; Figure S2F). There was no difference in the proportion of L3 RS, L5 RS, and L5 IB cells which expressed sADPs, nor the amplitude of sADPs in these cell types (Figure S2E & F). These results show that carbachol-induced sADPs, the hallmark of persistent firing, are widespread across RS and IB neurons in RSG but uniquely absent from LR neurons. This finding, together with the lack of persistent firing in LRs, shows that LR cells are likely to compute in a very different way in the presence of acetylcholine, when compared to all of their neighboring principal neuronal subtypes.

Since LR neurons are unique to the retrosplenial cortex, but RS cells are ubiquitous across the brain, we next compared responses to cholinergic activation across superficial layers of cortical regions neighboring RSG. We hypothesized that the lack of persistent firing will remain unique to LR cells, and that RS cells in other regions will fire persistently as did L3 RS cells in RSG.

### Pyramidal cells in layers 2/3 of neighboring cortical areas show persistent activity

To explore cholinergic activation of superficial layers in cortical regions neighboring RSG, we recorded from pyramidal cells in layers 2/3 of the adjacent RSD (which lacks LR cells; Brennan et al., 2020), as well as the anterior cingulate (ACC) and prelimbic cortex (PrL). All neurons examined in each region were physiologically RS cells, and 80-100% of these layer 2/3 cells in RSD, ACC and PrL showed persistent resposnes (Figure 3). Thus, LR cells are unique even among pyramidal cells in layers 2/3 of neighboring regions, in not exhibiting any persistent response to muscarinic agonists. We also consulted the expression of muscarinic receptor genes in the superficial excitatory neurons of those neighboring regions in the published 10x scRNA-seq dataset (Yao et al. 2021). We found that the Gi-Gq Balance in those areas resembles that of the L3 RS cells in RSG and is again starkly different from the LR neurons in RSG (data not shown). This further emphasizes the remarkably distinct biophysical and transcriptomic properties of LR cells.

**Figure 3.**
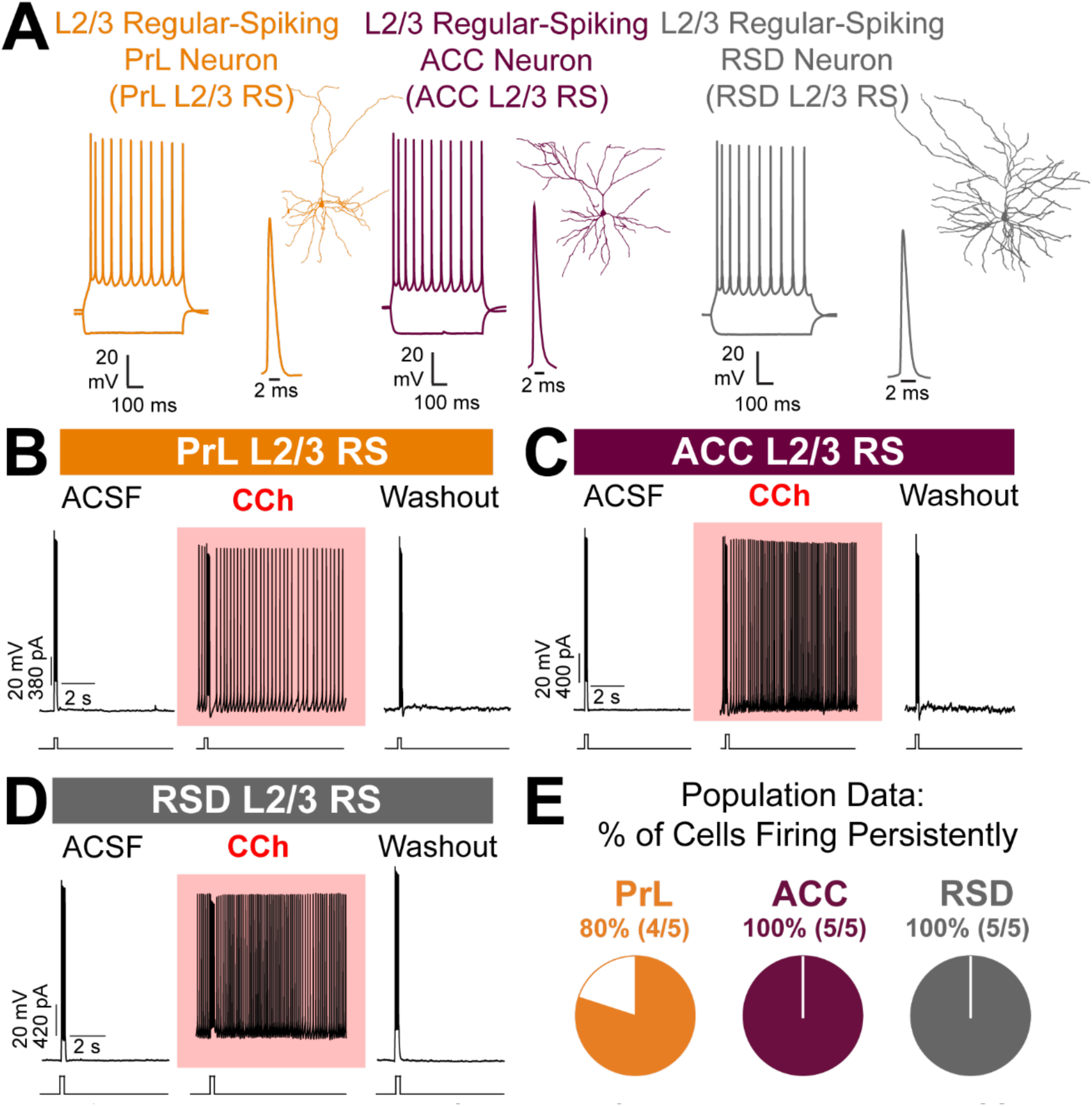
Excitatory neurons in superficial layers of cortical regions other than RSG show persistence. This is in stark contrast to superficial layers of RSG where LR cells, the dominant superficial excitatory cell type, do not fire persistently. ***A.*** Representative reconstructions and firing properties of L2/3 prelimbic cortex (PrL) neuron (orange; left), L2/3 anterior cingulate cortex (ACC, purple; middle), L2/3 dysgranular retrosplenial cortex (RSD, gray; right) neurons. All these cells show typical, regular-spiking (RS) firing patterns. ***B.*** Responses from a representative PrL neuron (shown in ***A***). Left: Baseline trace of the neuron’s activity in normal ACSF. Middle: Persistent firing after CCh. Right: Washout. ***C.*** Same as ***B*** but for a representative ACC neuron (shown in ***A***). ***D.*** Same as ***B*** but for a representative RSD neuron (shown in ***A***). ***E.*** Population analysis of the percentage of neurons in each region that exhibited persistent firing. The proportion of L2/3 neurons showing persistence was robustly high across the examined regions and did not significantly differ across regions.

Since carbachol is a non-selective muscarinic agonist, we next decided to investigate the mechanism underlying persistent firing to pinpoint which receptors carry this activity.

### Persistent firing in RSG is dependent on M1 receptor activity via an ICAN pathway

We used pharmacological blockers to determine the mechanism supporting persistent activity seen in RS and IB cells. Following the same protocol as previously described (Figure S1), layer 5 cells were exposed to CCh to induce persistent activity. Once the cell was firing persistently, we first tested whether persistent activity in RSG was dependent on glutamatergic or GABAergic synaptic activity by applying blockers— NMDA receptor antagonist APV, AMPA/kainite receptor antagonist CNQX, and GABA receptor antagonist picrotoxin—to the bath (see Methods). Application of these synaptic blockers did not affect persistent firing in any of the cells tested (0/5 cells blocked; Figure 4A), indicating that persistent activity in RSG does not rely on glutamatergic or GABAergic synaptic activity. We then verified that the CCh-induced persistent firing was indeed an effect of cholinergic signaling by adding the nonselective muscarinic receptor antagonist, atropine, to the bath. As expected, application of atropine blocked persistent firing (4/4; Figure 4B, top), indicating that persistence is an effect of muscarinic receptor activity induced by CCh. Using the M1-preferring antagonist, pirenzepine (Wess et al., 1991), we again observed a complete cessation of persistent firing in all tested neurons (4/4; Figure 4B, middle), indicating that persistent activity in RSG is dependent on M1 receptor activity, similar to previously described M1-dependent phasic and tonic responses in PrL neurons (Gulledge, et al., 2009; Dasari et al., 2017). Lastly, we applied flufenamic acid, a blocker of the calcium-activated nonselective cation current (*I*CAN) and found that persistent activity was blocked in the vast majority of cells tested (7/8; Figure 4B, bottom). These results are consistent with observations in other brain regions that exhibit persistent firing and suggest that persistent firing seen in RSG RS and IB cells is dependent on an M1- and ICAN-dependent pathway (Muñoz et al., 2017; Muñoz and Rudy, 2017; Navaroli et al., 2012; Yan et al., 2009).

**Figure 4.**
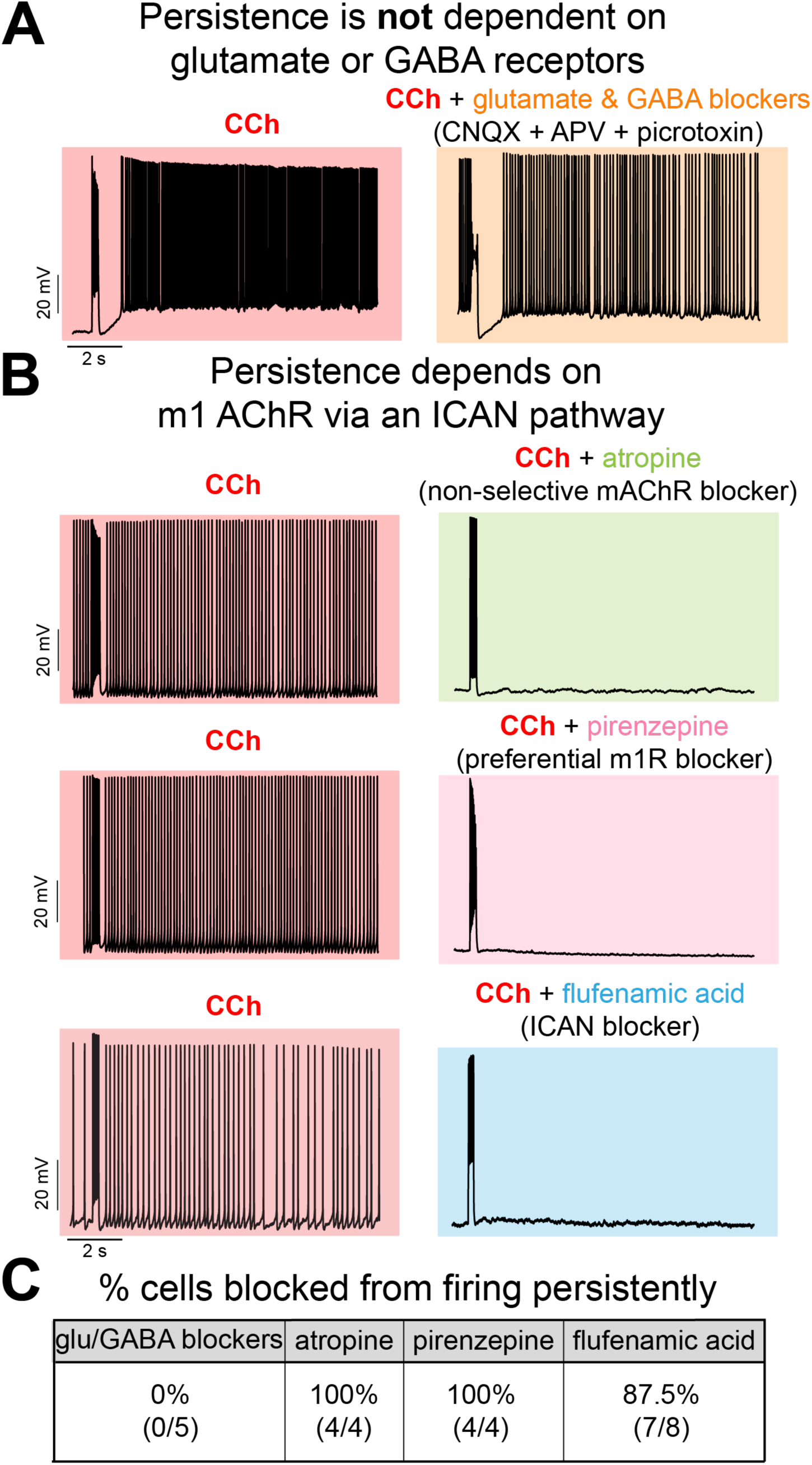
Persistence is independent of glutamatergic and GABAergic activity and dependent on the M1 receptor via an ICAN pathway. ***A.*** Left: Trace of persistent firing after exposure to CCh from a representative L5 RS neuron. Right: Trace from the same neuron after glutamatergic and GABAergic synaptic blockers APV, CNQX, and picrotoxin were added to the bath. Note that application of these glutamate and GABA blockers did not block persistent activity. ***B.*** *Top.* Same as ***A*** but for the nonselective muscarinic receptor antagonist, atropine. Application of atropine completely blocked persistent activity, revealing its dependence on muscarinic receptors. *Middle.* Same as ***A*** but for the preferential M1 receptor antagonist, pirenzepine. Application of pirenzepine completely blocked persistent activity, highlighting the predominant role of the M1 receptor in supporting persistent activity. *Bottom.* Same as ***A*** but for the ICAN blocker, flufenamic acid. Application of flufenamic acid blocked persistent activity, showing that the ICAN pathway is a necessary component. ***C.*** Summary of the percent of cells with persistent activity blocked by the pharmacological interventions reported in the previous panels. Persistent firing in dependent on M1 receptors, as well as the ICAN pathway.

Since layer 5 cells can show great morphological heterogeneity and the strength of cholinergic activation can differ across apical versus basal dendrites (Leung et al., 2010) we next examined whether morphological features of layer 5 cells correlated to their ability to show cholinergically-evoked persistent firing.

### Persistent activity is widespread across layer 5 cells, regardless of anatomical or morphological features

Principal cells of L5 RSG exhibit a variety of morphological features (Wyss et al., 1990, Yousuf et al., 2020). We therefore next investigated if the morphological properties of L5 pyramidal cells correlate with their ability to persistently fire (Hattox and Nelson, 2007; Tsiola et al., 2003; Wyss et al., 1990). To determine whether the somatic position and dendritic morphology of L5 RSG cells correlate with the ability to fire persistently, we reconstructed a subset of the tested layer 5 cells, including both L5 RS and L5 IB neurons (Figure S3A). We found that the depth of the neuron within layer 5, analyzed as distance of the soma from the L3/5 boundary, was independent of the cell’s likelihood to fire persistently (Figure S3B; chi-squared test, see Figure S3 legend for exact p-values). We next analyzed the dendritic morphology of the reconstructed layer 5 cells and found that proportion of persistently firing cells was independent of both apical and basal dendritic branching (Figure S3C & D; chi-squared test, see Figure S3 legend for exact p-values). We also examined potential sex differences in persistent activity and found that the proportion of cells which fired persistently did not significantly differ between male and female mice (chi-squared test, χ^2^(1, *n* = 45) = 1.38, p = 0.2393). Taken together, these results indicate that persistent activity is a pervasive feature of layer 5 pyramidal subtypes, regardless of somatic position, morphology, or sex.

### Lack of persistent activity in LR cells supports their role in encoding of angular head speed

The RSG has been predicted (Brennan et al., 2021), and experimentally confirmed (Keshavarzi et al., 2022), to play an important role in representing angular velocity information. Previous theoretical work suggests that thalamocortical synapses onto LR cells show short-term synaptic depression and these depressing synaptic dynamics can robustly encode angular head speed (Brennan et al., 2021). However, little is known about what would happen to this coding scheme in the presence of cholinergic agonists. This question is of importance given the large changes in levels of acetylcholine seen in the RSG during navigation versus standing still (Anzalone et al., 2009). We therefore used computational modeling to test whether the lack of persistent activity in LR cells may further support their hypothesized role in angular velocity encoding. We hypothesized that the artificial addition of persistent ICAN-dependent activity in LR would disrupt angular velocity coding. We built an integrate-and-fire model of an LR cell which incorporated ICAN (analogous to other excitatory cells in RSG, and contrary to the physiological observations in LR cells above). Even low values of ICAN conductances led to a slow ADP in this LR model cell, while higher values caused persistent firing (Figure 5A). We then added 3000 presynaptic thalamic HD inputs to the LR cell and investigated its activity in the presence and absence of ICAN (Figure 5B). In the absence of ICAN the model reproduced the results of the morphologically realistic model used by Brennan et.al. (2021) and we found that the LR cell reliably encoded angular head speed (Figure 5C, D). This encoding was, however, dramatically disrupted in the presence of ICAN, even at 3% of the maximum CAN current used here (Figure 5C, D). The maximum correlation of LR firing rate with angular head speed decreased rapidly with increasing ICAN values (Figure 5E, F). These results indicate that lack of persistent activity in LR cells is necessary for their role in encoding angular head speed. They also suggest that, as cholinergic levels change across brain states (e.g. during locomotion versus quiet wakefulness) (Anzalone et al., 2009; Goff et al., 2023, Kametani and Kawamura, 1990; Marrosu et al., 1995), LR cell coding principles will be minimally impacted by any degree of persistent responses.

**Figure 5.**
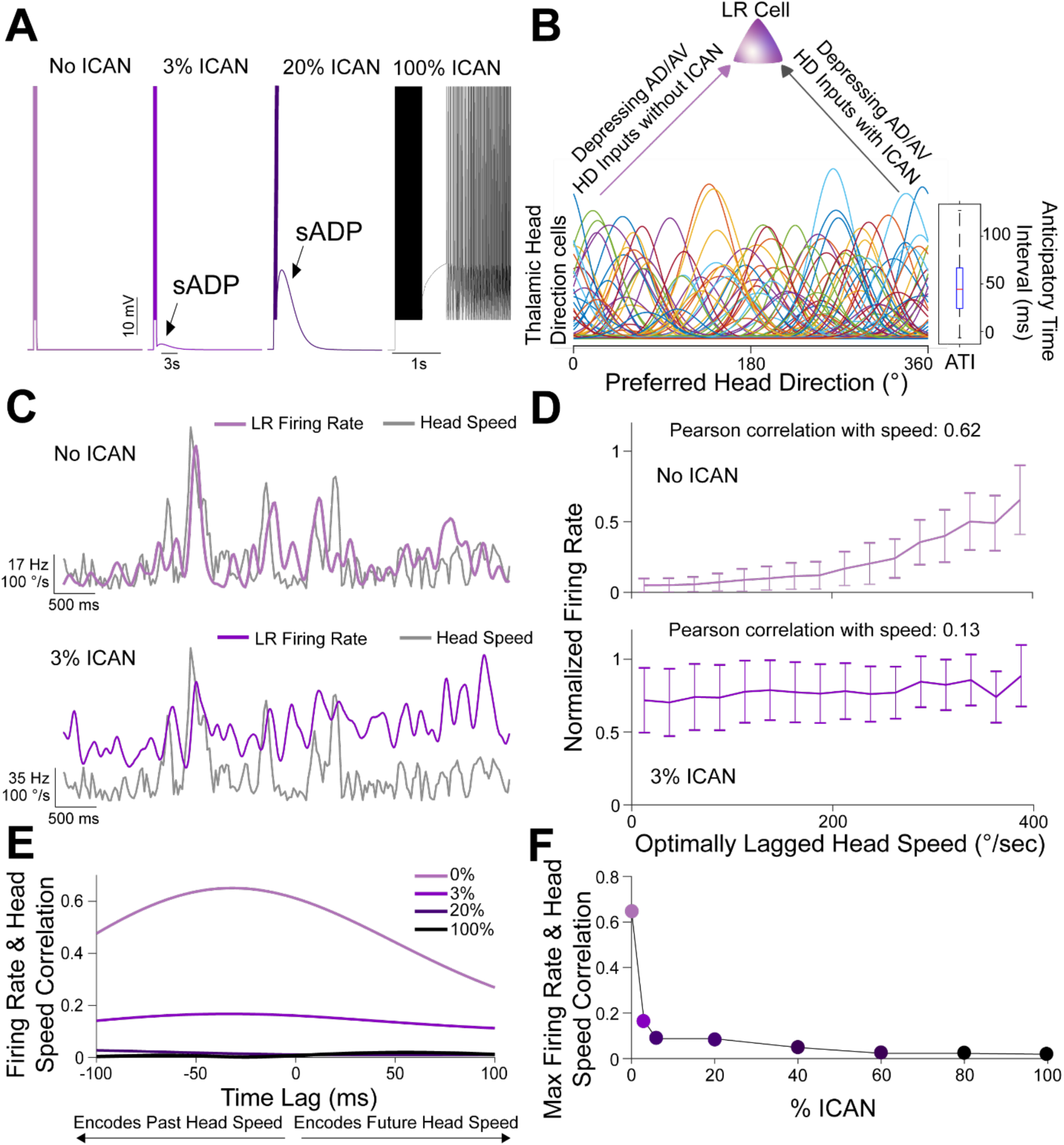
LR cells’ lack of cholinergically-evoked persistent firing allows them to rapidly encode angular head speed across cholinergic states. ***A.*** Responses from a model LR neuron. Left: Baseline response of the LR cell to a step current in the absence of a calcium-activated nonspecific cationic current (ICAN). This best matches the experimental observations of persistence in LR cells. Middle and Right Panels: Responses of the model to increasing ICAN current highlighting the slow ADP (sADP) and persistent firing. ***B.*** Schematic of the model with a heterogeneous population of 3000 HD cells providing input to a single LR neuron via depressing synapses with or without ICAN. Presynaptic HD cells had diverse tuning width, maximum firing rate, background firing rate, and anticipatory time interval (ATI). Tuning curves of a randomly selected subset of 100 HD cells shown here. Boxplot on the right depicts the distribution of ATIs of all presynaptic HD cells. ***C.*** Sample traces spanning five seconds of simulation time. Grey: Head turning speed; Light purple (top): Firing rate of LR cell in the absence of ICAN. Darker purple (bottom): Firing rate of LR cell with 3% ICAN. Note that the firing rate of an LR cell in the absence of ICAN visibly corresponds to the rapid fluctuations in head speed. In contrast, this faithful encoding is disrupted in the presence of ICAN. ***D.*** Firing rate of LR cell plotted against optimally-lagged head speed shows a strong correlation between speed and rate in the absence of ICAN (top). This encoding of speed is almost completely lost in the presence of even a weak ICAN (bottom). ***E.*** Cross-correlation between the model LR cell’s firing rate and head speed. The LR cell firing rate was maximally correlated at 31ms in the past but this encoding was almost completely disrupted with increasing values of ICAN added to the model cell. ***F.*** Maximum correlation of firing rate and head speed at different values of ICAN shows that the correlation decreases exponentially with increasing ICAN. Thus the encoding of angular head speed by LR cells is optimal without any ICAN, matching the experimentally observed lack of slow ADPs and persistent firing in LR cells.

Thus, our experimental findings highlight the unique suitability of LR neurons to encode angular head speed and suggest that this retrosplenial cell type may play a distinct computational role in RSG’s support of successful navigation independent of changes in cholinergic tone across brain states.

## DISCUSSION

### Retrosplenial Low Rheobase (LR) neurons represent a remarkably distinct pyramidal cell-type, highlighted by their complete lack of persistent firing in response to cholinergic activation and lack of M1-receptor expression

We examined the effects of cholinergic agonists on granular retrosplenial (RSG) pyramidal neurons at the cellular level. We found that regular-spiking (RS) and intrinsically-bursting (IB) cells across the superficial and deep layers fired persistently when exposed to the nonselective muscarinic receptor agonist carbachol (CCh). In layer 5 RS and IB cells, this persistence was independent of the cells’ physiological, morphological, anatomical, and sex characteristics, indicating that persistent activity is a widely shared feature of these neurons. Similar persistent responses to cholinergic agonists was seen in layer 2/3 cells of the RSD, ACC, and PrL. In contrast, we found that low-rheobase (LR) cells in L2/3 of the RSG, which have been identified as a key, selective recipient of directional and spatial inputs to RSG (Brennan et al., 2020, 2021), do not exhibit any persistent activity in response to cholinergic agonists. Persistent firing depends on muscarinic M1 receptors encoded by the Chrm1 gene (Figure 4). We found that LR cells express very low levels of Chrm1 transcripts (Figure 1), providing a molecular basis for the lack of cellular persistence in LR cells.

To the best of our knowledge, while the magnitude of sustained responses can vary within and across regions, persistent firing and/or sADPs have been found in all principal (glutamatergic) cortical cell types previously investigated. This includes principal cell types in deep layers of the primary visual, somatosensory, auditory and motor cortices, (Fu et al., 2019; Rahman and Berger, 2011), superficial and deep entorhinal cortex, including both stellate and pyramidal neurons (Magistretti et al., 2004; Jochems et al., 2013; Egorov et al., 2002; Tahvildari et al., 2007, Yoshida et al., 2018), superficial and deep subiculum (Kawasaki et al., 1999), and postsubiculum (Yoshida and Hasselmo, 2009), as well as superficial and deep cells of anterior cingulate, prelimbic, and perirhinal cortices (Baker et al., 2018; Dasari et al., 2017; Dembrow et al., 2010; Gulledge et al., 2009; Gulledge, 2024; Haj-Dahmane and Andrade, 1998; Lei et al., 2014; Navaroli et al., 2012; Ratté et al., 2018; Yan et al., 2009; Zhang and Seguela, 2010). Thus, LR neurons, unlike any other principal neurons reported, are unique in their complete lack of cholinergic-induced sADPs and persistent firing. What does this mean for the computations that LR cells can carry out?

### Implications for angular velocity encoding across brain states

From a computational perspective, the output of a neuron depends on two central factors: the inputs it receives and how its intrinsic properties process those inputs. So, what inputs do LR cells receive, how do they process them, and how does the lack of cholinergically-induced persistence impact this processing? We have previously shown that LR cells are selective recipients of spatial information from both the anterior thalamus and the dorsal subiculum (Brennan et al., 2021). The majority of neurons in the anterior thalamus encode head direction (Taube, 1995; Stackman & Taube, 1997; Taube & Muller, 1998; Ajabi et al., 2023) while neurons in the dorsal subiculum represent a diverse range of spatial and direction variables (Lever et al., 2009; Derdikman, 2009; Olson et al., 2017; Simonnet and Brecht, 2019; Bicanski and Burgess, 2020; Kitanishi et al., 2021). In RSG, RS cells near the layer 3/5 boundary receive inputs from so-called higher order regions such as the anterior cingulate and claustrum, as well as the contralateral RSG (Brennan et al., 2021). These higher order inputs largely ignore LR cells, highlighting the fact that LR cells are ideally positioned to process spatial inputs in a dedicated manner without the computational distraction from other inputs.

What do LR cells do with these spatial inputs? While the cellular processing of diverse subicular inputs remains an area of active exploration (Brennan et al., 2021; Kitanishi et al., 2021; Nitzan et al., 2020; Yamawaki 2019a), the processing of thalamic head direction inputs is more amenable to direct theoretical analysis. Thalamic synapses onto LR neurons show pronounced short-term depression (Brennan et al., 2021). Coupled with LR cells’ lack of intrinsic spike-frequency adaptation (Brennan et al., 2020) this depression of synaptic inputs allows LR neuron models to rapidly and faithfully compute the derivative of head direction: angular head velocity (AHV; Figure 5; Brennan et al., 2021). This represents an underappreciated way for cortical circuits to recompute AHV, a variable that is also calculated in mammalian brainstem circuits from vestibular, proprioceptive, and motor efference information (Bassett & Taube, 2001; Cullen & Taube, 2017; Hulse et al., 2020; Knierim & Zhang, 2012; Sharp et al., 2001). Cortical representations of AHV are likely to serve subtly different (more cognitively involved) computational functions than those served by brainstem AHV codes. The AHV calculated by RSG LR cells may be utilized by downstream retrosplenial layer 5 neurons and subsequently by entorhinal, RSD, and superior colliculus cells for path integration, egocentric boundary vector, and escape-to-shelter angle computations (Brennan et al., 2021; Alexander et al., 2020; Campagner et al., 2023; Ju & Gaussier, 2020; these retrosplenial computations are also fed back to the anterior thalamus to influence thalamic head direction coding (Clark et al., 2010) and cholinergic antagonists can disrupt the stability of anterior thalamic head-direction coding across environments (Yoder et al., 2017); see also Turner-Evans et al., 2017 for how angular velocity signals are computationally utilized in drosophila head direction circuits).

How does the lack of cholinergically-induced persistent firing impact LR coding of AHV? Cholinergic release in retrosplenial cortex ramps up dramatically during states of navigation-related locomotion (increased linear speed) compared to states with no linear motion (Anzalone et al., 2009), accompanied by precise changes in theta-coupled gamma and spline rhythms in RSG (Alexander et al., 2018; Ghosh et al., 2022). However, animals can rotate their heads when their bodies are otherwise still (zero linear speed, with low cholinergic levels) and also when locomoting (high linear speed, with high cholinergic levels; Anzalone et al., 2009). If LR cells had cholinergic M1- receptor-dependent persistent firing, they would not be able to rapidly compute the derivative of head direction, AHV, during the high cholinergic states associated with navigation (Figure 5). This is because the added input-independent depolarization from M1-dependent ICAN activation would render them far less sensitive to encoding rapid changes in synaptic input. Thus, while cholinergic release facilitates the persistent firing of cortical head direction neurons (Yoshida and Hasselmo, 2009, Taube et al., 1990; Cho and Sharp, 2001; Jacob et al., 2017), the same cholinergic activation would be deleterious to the rapid coding of angular head velocity (Figure 5). By lacking M1 receptors, LR neurons can implement a unique computational algorithm optimized for rapid, faithful encoding independent of cholinergic changes across navigational brain states.

More broadly, these results highlight the evolution of two parallel schemas for optimal spatial computation in retrosplenial circuits. RSG RS cells resemble more standard cortical pyramidal cells: they show spike-frequency adaptation (Brennan et al., 2020) and M1-dependent slow ADPs and persistence (Figure 2). Neighboring LR cells, only found in the RSG, are very different: they show no spike-frequency adaptation (Brennan et al., 2020) and no M1-dependent slow ADPs or persistence. These intrinsic differences are amplified by the aforementioned differences in synaptic input sources (Brennan et al., 2021). These parallel circuits are unlikely to remain completely segregated for long, with LR neurons providing the output of their computations to RS cells in RSG layer 5 (Brennan et al., 2021), as well as communicating via shared fast-spiking inhibitory interconnections (Brennan et al., 2020), ensuring that orientation and angular velocity information is combined for optimal RSG network computations. Thus, LR and RS cells are likely to serve complementary roles to support the computations needed for spatial orientation-related decision making, such as escape to shelter in fearful situations (Campagner et al., 2023, Keene & Bucci, 2008; Alexander et al., 2023). The remarkably distinct role of LR neurons in state-independent cortical computations is further supported by the fact that they are not synchronized by slow oscillations under urethane anesthesia even though their neighboring RS cells are locked to the slow rhythms (Mizuno & Ikegaya, 2024).

### Limitations of the study

Cholinergic neurotransmission in sensory cortical regions impacts thalamocortical signaling by modulating both pre- and postsynaptic signals (Arroyo et al., 2014, 2012; Kimura, 1999; Kruglikov & Rudy, 2008; Luchicchi et al., 2014; Muñoz et al., 2017; Muñoz and Rudy, 2017; Picciotto et al., 2012, Ferguson and Cardin, 2020; Minces et al., 2017). Acetylcholine can thus amplify sensory signals to increase the gain of relevant inputs over background signals or pre-established firing patterns (Donoghue and Carroll, 1987; Hasselmo & Giocomo, 2008; Kimura, 1999; Pinto et al., 2013; Polack et al., 2013; Soma et al., 2012, 2013; Picciotto et al., 2012). How acetylcholine impacts presynaptic thalamic terminals in RSG inputs remains unknown. Thus, a limitation of our study is that it does not focus on presynaptic effects of acetylcholine in RSG. Based on prior work in sensory cortices, we hypothesize that acetylcholine may slightly reduce the strength of thalamic inputs onto LR neurons, which may further help to maintain or amplify the LR neuronal AHV code during navigation by compensating for the small increases in firing rates of anterior thalamic head direction neurons seen during faster linear speeds (Taube & Bassett, 2003) and perhaps also relate to representational drift seen in these thalamic head direction neurons (Ajiba et al., 2023). Future work will thus focus on how cholinergic agonists impact presynaptic thalamic terminals in RSG and how this further refines orientation and AHV coding in the RSG.

## ACKNOWLEDGMENTS

This work was supported by NIH R34NS127101 (OJA); NIH P50NS123067 (OJA); Alzheimer’s Association Grant AARG-NTF-21-846572 (OJA); NIH T32-DC000011 (CRK, TGE); NIH T32-DA007268 (TGE); NIH T32-NS076401 (EKWB, CRK); NSF graduate fellowship (EKWB).

## MATERIALS AND METHODS

### Animals

All animal use and housing were approved by the University of Michigan Institutional Animal Care and Use Committee. The following mice – all maintained on C57Bl6 backgrounds – were used in this study: Ai32 (Jackson Laboratories, 024109), *Camk2α*^Cre^ (Jackson Laboratories, 005359), *Camk2α*^Cre^ x Ai32 (crossed in house), C57Bl6-wildtype (Charles River, stock #027), Ai14 (Jackson Laboratories, 007914), *Grp*-KH288^Cre^ (RRID:MMRRC 037585-UCD), *Pvalb*^Cre^ (PV-IRES-Cre; Jackson Laboratories, 008069), *Pvalb*^Cre^ x Ai32 (crossed in house), Tg(Sla-cre)KJ303 (MGI:4940648), Tg(Sla-cre)KJ319 (MGI:4847420) x Ai32 (crossed in house), and *Grp*-KH288^Cre^ x Ai32 (crossed in house). LR cells showed no persistent activity in any of the mouse lines in which they were recorded.

### Tissue dissection and processing for 10x snRNA-seq

Mice were deeply anesthetized using isoflurane and decapitated. The brain was rapidly removed into ice cold ACSF (same composition as for patch clamp recordings, see below) and then placed in a brain matrix.1 mm thick coronal sections were cut with a razor blade at distance between -1.7 to -2.9 mm from bregma. Each slice was transferred to a glass culture dish filled with fresh ice cold ACSF. The slice was then cut in half width-wise just below the hippocampus and the lower half of the slice was discarded. To help limit the dissection to only the RSG region, hippocampus was first pulled off the top half of the slice, leaving only a strip of cortex, which included the area of interest. Then, a cut was made along the boundary of the RSG/RSD border in each hemisphere, and the dissected RSG from both hemispheres was combined and immediately flash-frozen in a PCR tube. Remaining cortical regions from the same strips were also saved to serve as controls for cell sorting. All tubes were kept at -80C until further processing.

For the subsequent nuclei isolation, microdissected RSG regions from 3 littermates were combined into a single sample and processed based on existing protocol (Affinati et al, 2021). Briefly, the combined tissue samples were homogenized in a lysis buffer using a glass dounce, filtered through a 30µm MAC strainer, and centrifuged at 500rcf for 5 min and the pellet resuspended in a wash buffer. Nuclei were filtered and strained again and resuspended in a wash buffer with propidium iodide and went through FACS sorting on a MoFlo Astrios Cell Sorter. Afterwards, the sample was centrifuged at 100rcf for 6 min and the pellet resuspended in a wash buffer for a concentration of 750–1200 nuclei/μL. The sample was then submitted for snRNASeq.

### Slice electrophysiology

#### Slice preparation

Detailed description of slice preparation is available elsewhere (Brennan et al, 2020). Briefly, mice were deeply anesthetized using isoflurane and decapitated. The brain was rapidly removed and placed in an ice cold, carbogen-saturated high sucrose solution. 300 µm thick slices were cut using Leica 1200VT or 1000S vibratomes and transferred into high magnesium artificial cerebrospinal fluid (ACSF) kept at 32 °C. After 20 min, the recovery chamber containing the slices was moved to room temperature. Experimental recordings started 45min after the additional room temperature recovery.

#### Whole-cell recordings

Slices were placed in a submerged recording chamber continuously perfused at 2 mL/min with body temperature ACSF containing: 126 mM NaCl, 1.25 mM NaH2PO4, 26 mM NaHCO3, 3 mM KCl, 10 mM dextrose, 1.20 mM CaCl2, and 1 mM MgSO4. Cells were visualized using an Olympus BX51WI microscope with Olympus 60x water immersion lens and Andor Neo sCMOS camera (Oxford Instruments, Abingdon, Oxfordshire, UK). Patch pipettes were pulled using a horizontal puller and had resistances between 4-7 MΩ. Recordings were done using a potassium gluconate intracellular solution containing: 130 mM K-gluconate, 2 mM NaCl, 4 mM KCl, 10 mM HEPES, 0.2 mM EGTA, 0.3 mM GTP-Tris, 14 mM phosphocreatine-Tris, and 4 mM ATP-Mg (pH 7.25, osmolarity 290 mOsm).

Patch clamp recordings were obtained with Multiclamp 700B and Digidata 1440A (Molecular Devices). Neurons from layers 2, 3, and 5 of RSG and L2 and 3 of other cortical regions were randomly chosen based on their shape, and recordings were done in current clamp mode. Only the cells with resting membrane potential of -55 mV or lower were used for the analysis. Cells were adjusted for series resistances and held at -65 mV for all but the persistence protocol recordings. Recordings were not liquid junction potential corrected. Each recorded cell was first characterized for intrinsic and firing properties (see Brennan et al., 2020, 2021 for detailed protocols) and subsequently tested for persistence.

#### Pharmacology

After the cells were characterized (see Methods above), the persistence protocol was started (see Figure S1 for schematic). The protocol consisted of repetitive sweeps of 3 current injections, each at an intensity at least twice that of the rheobase, meant to evoke persistent firing. These current injections were followed by a small negative step (same intensity and length as for input resistance characterization; see Brennan et al., 2020, 2021) to monitor the input resistance changes. From that point until the end of the experiment, the resting membrane potential was not adjusted. Cells were recorded for about 2 minutes in ACSF before the non-selective muscarinic receptor agonist, carbachol (40 uM; CCh), was added to the bath, and responses were recorded for 10 min. CCh was then washed out using standard ACSF for 5 min, or until no persistent activity was observed for at least 1 min.

In order to evaluate the nature of persistent firing, blockers were added to the recording chamber. AP-5 (50 µM), CNQX (10 µM), flufenamic acid (100 uM) and pirenzepine (1 µM) were added from frozen stock solutions, while picrotoxin (100 µM) and atropine (1 µM) were added fresh.

### Quantification and statistical analysis

#### snRNA-seq data analysis

RNA sequencing data was processed by the University of Michigan Advanced Genomics Core into a UMI counts matrix with 9,538 cells and 55,415 genes. For quality control, we removed all cells with fewer than 700 unique genes detected. We used the Python package Scrublet for doublet detection and removed all cells with a doublet score greater than 0.3. 8,447 cells remained after these filters were applied.

Clustering analysis was performed using Seurat’s FindClusters function, which identified 28 distinct cell clusters in our dataset. To interpret these clusters, we used Seurat’s FindTransferAnchors function for CCA analysis, mapping our cells onto clusters from the 10x single-cell RNA-seq data in Yao et al. 2021. The Yao et al. dataset consists of 1.1 million cells, but cluster mapping was performed only onto the 65,710 cells with the region label “RSP” (corresponding to the retroslenial cortex dysgranular and granular regions). From this analysis, we assigned names to each of our clusters depending on putative layer (when applicable) and cell type. These clusters included L2/3 Cxcl14, L2/3 Calb1, L5 IT, L6 IT, L5 PT, L6 CT, L5/6 NP, L6b, L5 Scnn1a, several inhibitory clusters, and some non-neuronal clusters. Two of our clusters were specifically excluded from downstream analysis: One consisted of 191 cells with high mitochondrial gene counts but low overall UMI counts, and the other of 25 cells with extremely high UMI counts compared to other clusters.

Python’s “umap-learn” package was used to generate the UMAP in figure 1. We first narrowed our dataset down to 10 putative excitatory neuronal clusters, totaling 3,606 cells. From these cells, we identified 2000 highly variable genes to use as inputs to the UMAP algorithm by computing each gene’s normalized variance, defined as variance divided by mean UMI counts. Scran and Seurat were used to normalize the UMI data for these 2000 genes. The resulting matrix was used as the input to principal component analysis (PCA), from which 25 principal components (PCs) were generated. These PCs served as inputs for the UMAP algorithm.

Clusters were grouped in the Figure 1 UMAP as follows: L2/3 Cxcl14 and L2/3 Calb1 were each in their own groups; L5 IT, L5 PT, and L5 Scnn1a were assigned “L5”; and L6 CT, L5/6 NP, and L6b were assigned “L6”.

#### Electrophysiological measurements and analysis

Cells were characterized based on their spike threshold, spike amplitude, spike width, spike frequency adaptation ratio, latency to first spike, rheobase, input resistance (Rin), input capacitance (Cin), and membrane time constant (τm), as described elsewhere (Brennan et al., 2021), and all measurements were obtained using Clampfit analysis and MATLAB scripts.

Distance from layer 3 was measured in µm using Andor Solaris software and defined as the distance between the boundary between layers 3 and 5 and the tip of the recording pipette.

Slow afterdepolarization (sADP) was measured in Clampfit as the highest amplitude of the membrane potential within 1.5 s after the end of the current injection, relative to 0.5 s baseline before the current step for three consecutive steps, and the average of those steps was reported. Initial sADP was measured from the first three step currents in the persistence protocol, while onset sADP was measured from 3 steps immediately preceding persistent firing. For 3 cells’ measurements, those time points were slightly offset due to either nonstandard protocol (1 cell) or EPSPs being too high to reliably measure the sADP at the typical time points (2 cells). In those cases, first reliable step currents immediately adjacent to the standard ones were used. For cells that did not fire persistently, CCh sADP was measured from three consecutive current steps starting 5 min after carbachol entered the bath. If no sADP was observed, the sADP was reported as 0 mV.

Neurons were classified as persistently firing if they exhibited at least 6 spikes within 13 s after the termination of the current injection (modified from Jochems et al., 2013, to account of the different current step interval) in at least two of three consecutive step currents.

### Morphological reconstructions

Biocytin (5 mg/ml) was added to the intracellular recording solution to aid morphological analysis. During whole-cell recordings, biocytin diffused into cells for at least 30 min. At the end of each recording, an additional zap protocol (15 1-4 nA current injections at 1 Hz) was applied to improve the diffusion process (Jiang et al., 2015). After the recording ended, the patch pipette was carefully withdrawn from the cell and the slice was left to rest in the recording chamber for 10 min before being transferred to 4% PFA solution for 24-72 h fixation. After fixation, the slices were washed in phosphate buffer solution (PBS) and incubated in PBS containing 0.4% triton and 1:1000 streptavidin conjugated Alexa Fluor 647 for another 48 h. Slices were then again washed in PBS and mounted on slides using FluoromountG mounting medium and left to set for at least 24 h before imaging.

Fluorescent fills were imaged with Zeiss Axio Image M2 confocal microscope with 20x lens as z-stacks with z-step of 0.5 µm. Neurons were then reconstructed from these files using NeuTube software.

#### Morphological analysis

17 (13 firing persistently and 4 not) reconstructed neurons (in the .swc format) were quantified using the NeuroM Python module. The number of basal dendrite terminations was counted using Python scripts, and the number of apical tuft terminations was counted by hand, to avoid including oblique dendrites.

### Computational modeling

An integrate and fire model was used to simulate the activity of LR cell. The dynamics of the membrane potential is given by

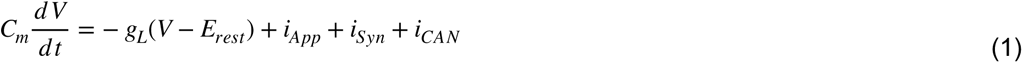

where V is the membrane potential of the cell and *Cm* denotes the capacitance of the membrane (1 μF/cm^2^). *Erest* is the resting membrane potential (-70 mV). *gL* is the conductance of the leak channels (0.1 mS/cm^2^). *Isyn* is the synaptic current to the cell and *ICAN* is the calcium-activated nonspecific (CAN) cationic current. i*App* refers to the spontaneous firing of the cell and is modelled as an external Poisson input. Each time the membrane potential exceeds the threshold voltage (-47mV), and action potential is recorded and the voltage is reset to -52mV.

### Synaptic Current

The presynaptic activity of 3000 HD cells is modelled similar to Brennan et al., 2021. Briefly, each HD cell has a Gaussian tuning curve :

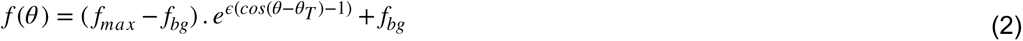

where fbg is the background firing rate, fmax is the maximum firing rate, and ɛ sets the tuning curve width and θ is the preferred angle for that cell. θT(t)∈[0,2π] denotes the head direction at time t ms. The synaptic current to the LR cell is given by

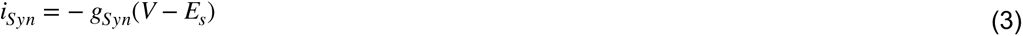

where Es is the reversal potential of the synapse (20mV). Each synapse is defined by a set of two parameters -

x(t): the proportion of vesicles available at time t.

u(t): the release probability at time t

Synaptic depression is modelled as a depletion of synaptic vesicles in response to each pre-synaptic spike followed by slow recovery of the vesicle pool (τr = 323ms) with a minimum fraction of x (U = 0.0958) contributing to the conductance on each presynaptic spike. The release probability increases transiently with each presynaptic spike and decays with time constant τf (30ms).

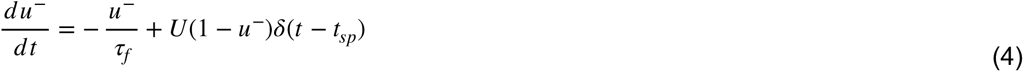

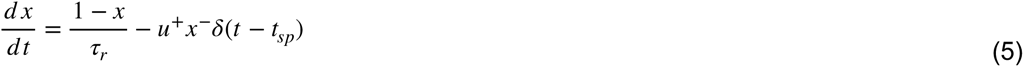

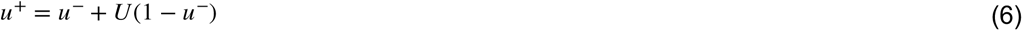

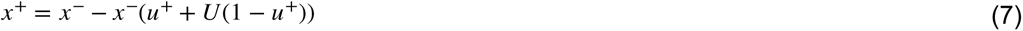

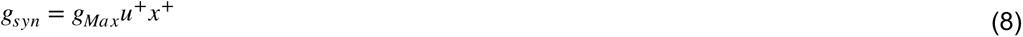

where gMax (0.0082 mho) is the maximum synaptic conductance.

### ICAN Current

The ICAN model was adopted from Liu et al. (2019) and follows the dynamics given below

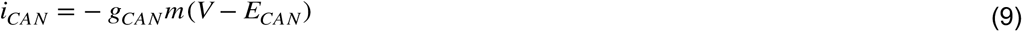

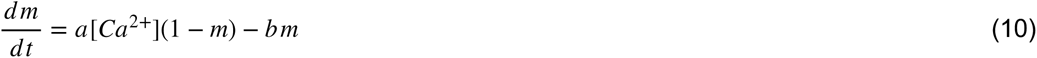

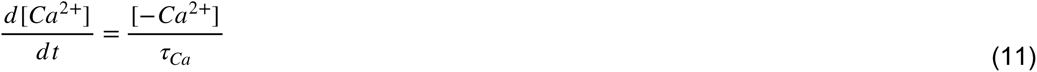

*E*CAN (-20mV) is the reversal potential of the CAN current ion channels and *m* is a dimensionless quantity between 0 and 1 associated with the activation of the CAN current ion channels. *a* and *b* are free parameters and were chosen to be 0.01 and 1 respectively. Each post-synaptic spike leads to an influx of a fixed amount of calcium (kCa = 5e-4) and the current decays with a time constant (τCa) of 1000ms. gCAN is the maximum ion conductance and was varied from 0.01029 to 0.3 mho. Ca^2+^ is the time varying calcium ion concentration.

---

**Figure S1.**
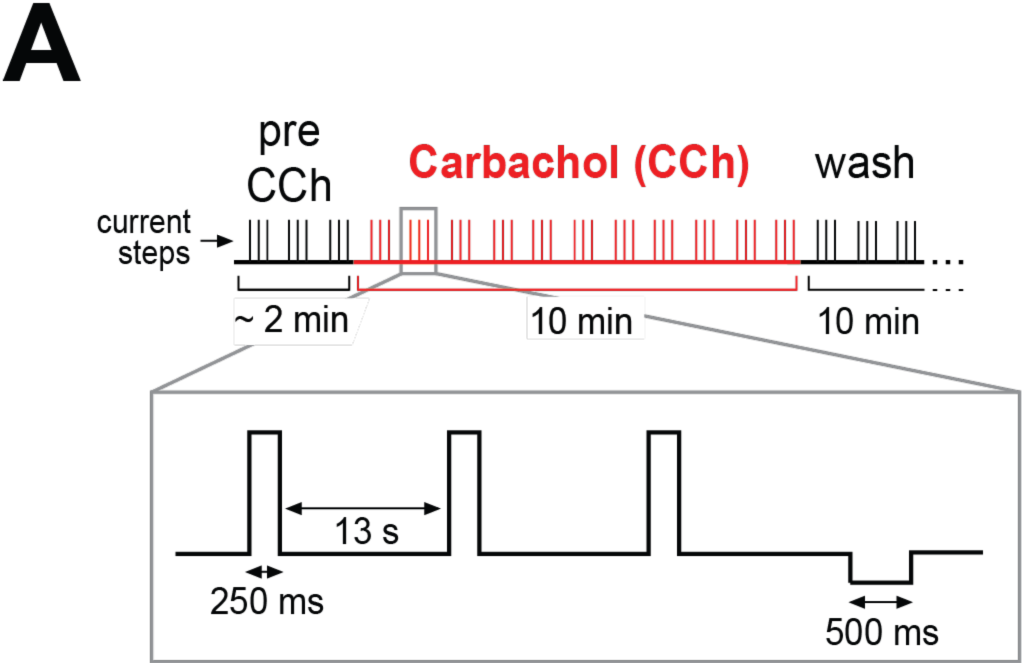
Schematic of the full recording protocol with an inset showing a single sweep. See Methods for details.

**Figure S2.**
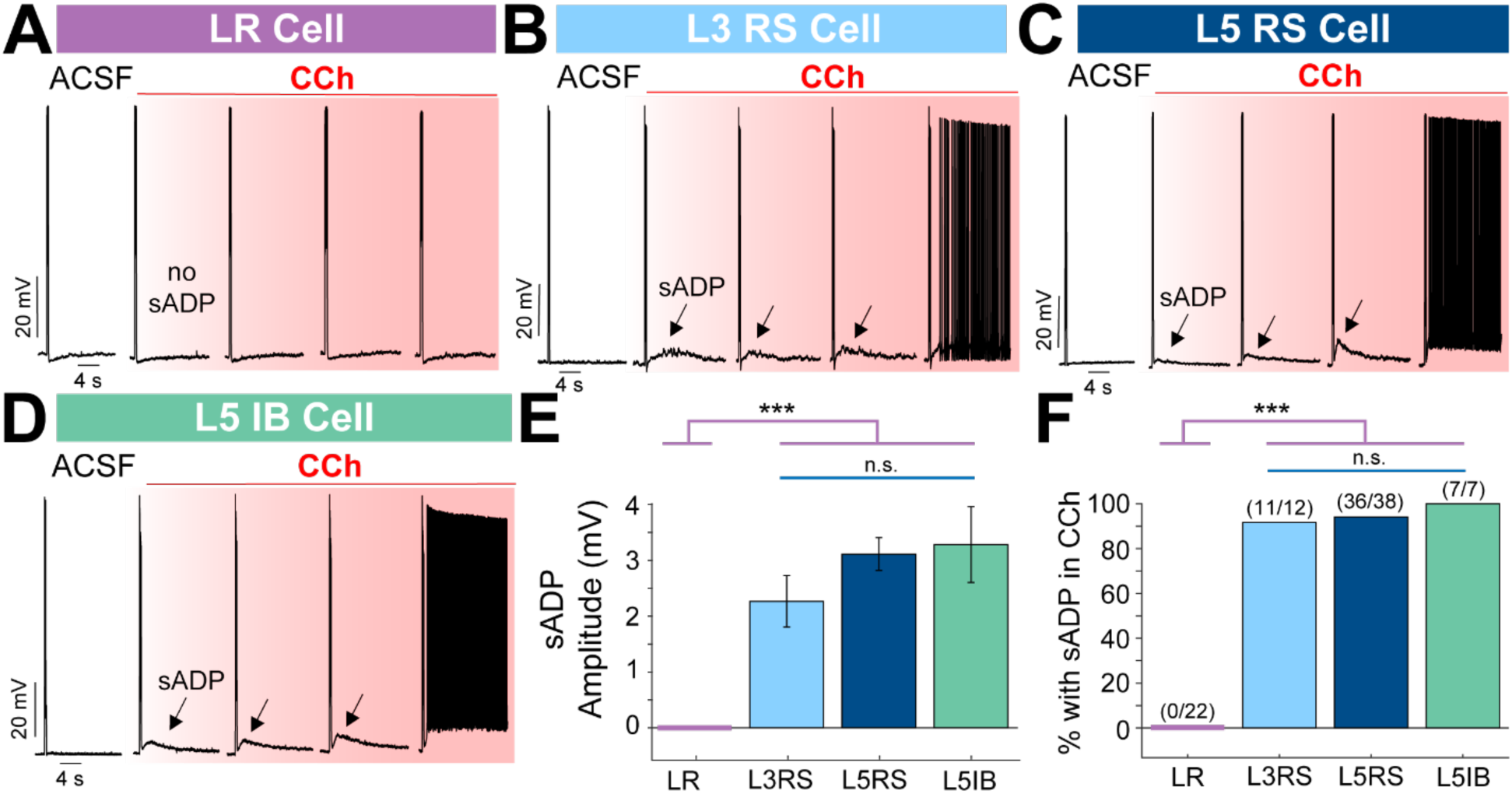
Slow ADP is a precursor of persistent firing and completely absent in LR neurons. ***A.*** Responses from a representative LR neuron. Left-most trace: baseline response to current injection, trace of LR activity in normal ACSF. Further traces: responses to the same current injection protocol in the presence of CCh at progressively increasing time points. Note that LRs show complete lack of sADP response to cholinergic activation. ***B.*** Same as **A**, but responses from a representative L3 RS neuron. Note the slow build-up of sADP that immediately preceded persistent firing (sweep not shown). ***C.*** Same as ***B*** but for a L5 RS neuron. Build-up of slow ADP leading to persistent firing is again clearly visible. ***D.*** Same as ***B*** but for a L5 IB. This cell type also shows robust sADP build-up in response to CCh. ***E.*** Population analysis of sADP amplitudes across the four cell types. LR neurons had significantly smaller sADPs (0 mV) compared to all other cell types (LR v L3 RS: p 1.74e-07; LR v L5 RS: p = 3.60e-10; LR v L5 IB: p = 3.24e-07; rank sum test). In contrast, sADP amplitude did not significantly differ between L3 RS, L5 RS, or L5 IB cells (L3 RS v L5 RS: p = 0.1915; L3 RS v L5 IB: p = 0.1025; L5 RS v L5 IB: p = 0.8777; rank sum test). ***F.*** Population analysis of sADP occurrence across the four cell types. LR neurons had a significantly lower percentage of cells exhibiting sADPS (0%) compared to the other cell types (LR v L3 RS: Χ^2^(1, *n* = 36) = 29.812, p = 4.76E-08; LR v L5 RS: Χ^2^(1, *n =* 60) = 52.105, p = 5.26E-13; LR v L5 IB: Χ^2^(1, *n* = 29) = 29.00, p = 7.24E-08; chi-squared test for all). In contrast, percentage of cells exhibiting sADPs did not differ between L3 RS, L5 RS, or L5 IB cells (L3 RS v L5 RS: Χ^2^(1, *n* = 50) = 0.15, p = 0.6962; L3 RS v L5 IB: Χ^2^(1, *n =* 19) = 0.616, p = 0.4326; L5 RS v L5 IB: Χ^2^(1, *n* = 45) = 0.386, p = 0.5346; chi-squared test for all).

**Figure S3.**
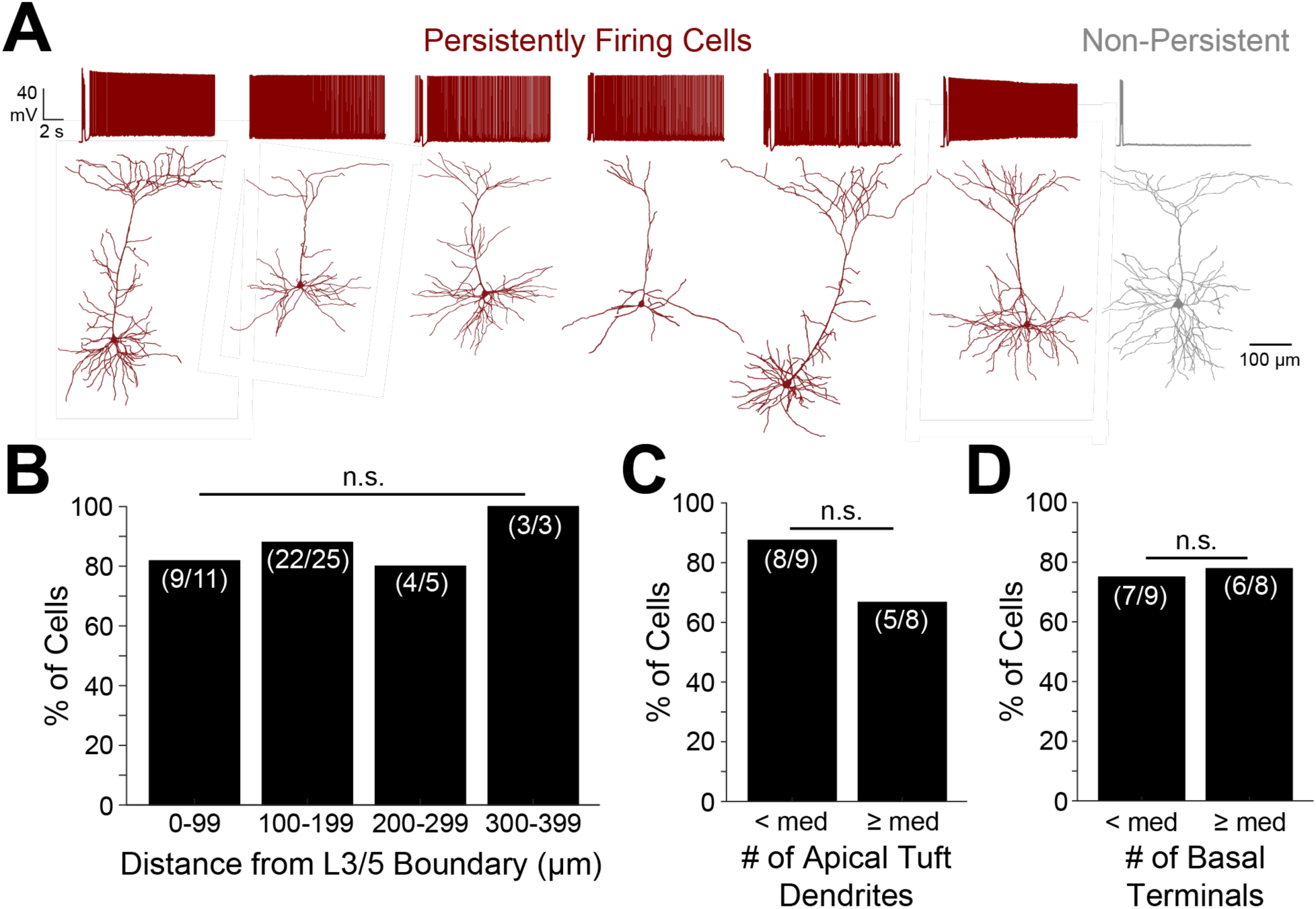
Persistent activity in layer 5 RSG neurons is independent of anatomical and morphological features. ***A.*** Representative reconstructions of a subset of layer 5 neurons which either did (red) or did not (grey) fire persistently. Traces of the shown cell’s response in CCh are shown above each reconstruction. ***B.*** Population analysis of proportion of recorded L5 cells which fired persistently plotted against soma distance from the L3/5 boundary (n=44, one cell did not have an image to measure the distance). There is no significant difference in propensity to fire persistently across the cortical depth (0-99 v 100-199: Χ^2^(1, *n* = 36) = 0.24, p = 0.6213; 0-99 v 200-299: Χ^2^(1, *n* = 16) = 0.01, p = 0.9312; 0-99 v 300-399: Χ^2^(1, *n* = 14) = 0.64, p = 0.4250; 100-199 v 200-299: Χ^2^(1, *n* = 30) = 0.23, p = 0.6310; 100-199 v 300-399: Χ^2^(1, *n* = 28) = 0.40, p = 0.5254; 200-299 v 300-399: Χ^2^(1, *n* = 8) = 0.69, p = 0.4076; chi-squared test for all). ***C.*** Population analysis of proportion of reconstructed cells which fired persistently plotted against number of apical tuft dendrites with bins split by median. There is no significant difference in propensity to fire persistently across the two groups (Χ^2^(1, *n* = 17) = 1.02, p = 0.3121; chi-squared test). ***D.*** Same as ***C*** but for number of basal terminals. There is no significant difference in propensity to fire persistently across the two groups (Χ^2^(1, *n* = 17) = 0.02, p = 0.8928; chi-squared test).

**Table 1.** Intrinsic cell properties of LR, L3 RS, L5 RS, and L5 IB neurons. LRs significantly differed from L3RS cells on: p<0.001 – input resistance, input capacitance, adaptation ratio, spike width, spike amplitude, and latency to first spike; p<0.01 – spike threshold and rheobase; LRs significantly differed from L5RS cells on: p<0.001 – input resistance, input capacitance, adaptation ratio, spike amplitude, spike threshold, latency to first spike, and rheobase; p<0.01 – membrane time constant. LRs significantly differed from IB cells on: p<0.001 – input capacitance and latency to first spike; p<0.01 – adaptation ratio and input resistance; p<0.05 – spike width, and spike amplitude. L3RS cells significantly differed from L5 RS cells on: p<0.001 – membrane time constant, spike width, and rheobase; p<0.01 – input resistance; p<0.05 – spike amplitude. L3RS cells significantly differed from IB cells on: p<0.05 – spike amplitude. L5RS cells significantly differed from IB cells on: p<0.01 – spike width; p<0.05 – membrane time constant and spike amplitude.

|  | LR (n=22) | L3 RS (n=12) | L5 RS (n=38) | L5 IB (n=7) |
| --- | --- | --- | --- | --- |
| Postnatal age at time of recording (days) | 42.95 ± 3.49 | 30.75 ± 1.11 | 31.63 ± 1.0 | 36.0 ± 4.36 |
| Resting potential (mV) | -66.09 ± 0.91 | -63.0 ± 1.46 | -61.37 ± 0.53 | -62.71 ± 1.43 |
| Input resistance (MΩ) | 540.28 ± 29.98 | 171.73 ± 21.89 | 106.31 ± 11.38 | 234.54 ± 60.95 |
| Input capacitance (pF) | 34.52 ± 1.33 | 125.22 ± 11.90 | 144.54 ± 8.01 | 114.09 ± 23.0 |
| Membrane time constant (ms) | 17.96 ± 0.55 | 19.27 ± 1.55 | 12.95 ± 0.65 | 18.88 ± 2.83 |
| Action potential threshold (mV) | -39.46 ± 0.43 | -43.53 ± 0.76 | -43.03 ± 0.42 | -40.95 ± 1.07 |
| Action potential amplitude (mV) | 68.68 ± 2.04 | 86.99 ± 2.09 | 80.13 ± 1.45 | 73.05 ± 3.49 |
| Action potential width (ms) | 0.8 ± 0.06 | 1.11 ± 0.05 | 0.71 ± 0.03 | 1.20 ± 0.21 |
| Spike frequency adaptation ratio | 1.08 ± 0.04 | 2.0 ± 0.13 | 4.0 ± 0.90 (n=37) | 9.32 ± 3.37 |
| Latency to first spike (ms) | 834.26 ± 25.07 | 330.09 ± 64.51 | 375.05 ± 47.14 | 316.24 ± 57.07 |
| Rheobase (pA) | 31.54 ± 1.87 | 73.62 ± 16.30 | 132.60 ± 12.79 | 90.77 ± 34.24 |

## REFERENCES

1. Affinati AH, Sabatini PV, True C, et al. 2021. Cross-species analysis defines the conservation of anatomically segregated VMH neuron populations. Elife 10:e69065. doi:10.7554/eLife.69065

2. Ajabi Z, Keinath AT, Wei XX, Brandon MP. Population dynamics of head-direction neurons during drift and reorientation. Nature. 2023 Mar;615(7954):892-899. doi: 10.1038/s41586-023-05813-2. Epub 2023 Mar 22. Erratum in: Nature. 2023 May;617(7961):E10.

3. Alexander AS, Carstensen LC, Hinman JR, Raudies F, Chapman GW, Hasselmo ME. Egocentric boundary vector tuning of the retrosplenial cortex. Sci Adv. 2020 Feb 21;6(8):eaaz2322. doi: 10.1126/sciadv.aaz2322.

4. Alexander AS, Nitz DA. Retrosplenial cortex maps the conjunction of internal and external spaces. Nat Neurosci. 2015;18(8):1143–1151. doi:10.1038/nn.4058

5. Alexander AS, Nitz DA. Spatially Periodic Activation Patterns of Retrosplenial Cortex Encode Route Sub-spaces and Distance Traveled. Curr Biol. 2017;27(11):1551–1560.e4. doi:10.1016/j.cub.2017.04.036

6. Alexander AS, Place R, Starrett MJ, Chrastil ER, Nitz DA. Rethinking retrosplenial cortex: Perspectives and predictions. Neuron. 2023 Jan 18;111(2):150–175. doi: 10.1016/j.neuron.2022.11.006.

7. Alexander AS, Rangel LM, Tingley D, Nitz DA. Neurophysiological signatures of temporal coordination between retrosplenial cortex and the hippocampal formation. Behav Neurosci. 2018 Oct;132(5):453–468. doi: 10.1037/bne0000254.

8. Andrade R. 1991. Cell excitation enhances muscarinic cholinergic responses in rat association cortex. Brain Res 548:81–93. doi:10.1016/0006-8993(91)91109-E

9. Anzalone S, Roland J, Vogt B, Savage L. 2009. Acetylcholine efflux from retrosplenial areas and hippocampal sectors during maze exploration. Behav Brain Res 201:272–278. doi:10.1016/j.bbr.2009.02.023

10. Arnold HM, Burk JA, Hodgson EM, Sarter M, Bruno JP. 2002. Differential cortical acetylcholine release in rats performing a sustained attention task versus behavioral control tasks that do not explicitly tax attention. Neuroscience 114:451–460. doi:10.1016/S0306-4522(02)00292-0

11. Arroyo S, Bennett C, Aziz D, Brown SP, Hestrin S. 2012. Prolonged disynaptic inhibition in the cortex mediated by slow, non-α7 nicotinic excitation of a specific subset of cortical interneurons. J Neurosci 32:3859–3864. doi:10.1523/JNEUROSCI.0115-12.2012

12. Arroyo S, Bennett C, Hestrin S. 2014. Nicotinic modulation of cortical circuits. Front Neural Circuits. doi:10.3389/fncir.2014.00030

13. Baker AL, O’Toole RJ, Gulledge AT. 2018. Preferential cholinergic excitation of corticopontine neurons. J Physiol. 596(**9**):1659–1679. doi: 10.1113/JP275194.

14. Bassett JP, Taube JS. Neural correlates for angular head velocity in the rat dorsal tegmental nucleus. J. Neurosci. 21, 5740–5751 (2001).

15. Beierlein M, Gibson JR, Connors BW. A network of electrically coupled interneurons drives synchronized inhibition in neocortex. Nat Neurosci. 2000 Sep;3(9):904–10. doi: 10.1038/78809. PMID: 10966621

16. Berger-Sweeney J, Heckers S, Mesulam MM, Wiley RG, Lappi DA, Sharma M. 1994. Differential effects on spatial navigation of immunotoxin-induced cholinergic lesions of the medial septal area and nucleus basalis magnocellularis. J Neurosci 14:4507– 4519. doi:10.1523/jneurosci.14-07-04507.1994

17. Bianchi R, Wong RKS. 1994. Carbachol-induced synchronized rhythmic bursts in CA3 neurons of guinea pig hippocampus in vitro. J Neurophysiol 72:131–138. doi:10.1152/jn.1994.72.1.131

18. Bicanski A, Burgess N. Neuronal vector coding in spatial cognition. Nat Rev Neurosci. 2020 Sep;21(9):453–470. doi: 10.1038/s41583-020-0336-9.

19. Bigl V, Woolf NJ, Butcher LL. 1982. Cholinergic projections from the basal forebrain to frontal, parietal, temporal, occipital, and cingulate cortices: A combined fluorescent tracer and acetylcholinesterase analysis. Brain Res Bull 8:727–749. doi:10.1016/0361-9230(82)90101-0

20. Brennan EKW, Jedrasiak-Cape I, Kailasa S, Rice SP, Kumar Sudhakar S, Ahmed OJ. 2021. Thalamus and claustrum control parallel layer 1 circuits in retrosplenial cortex. bioRxiv. doi:10.1101/2020.09.17.300863

21. Brennan EKW, Sudhakar SK, Jedrasiak-Cape I, John TT, Ahmed OJ. 2020. Hyperexcitable Neurons Enable Precise and Persistent Information Encoding in the Superficial Retrosplenial Cortex. CellReports 30:1598–1612.e8. doi:10.1016/j.celrep.2019.12.093

22. Campagner D, Vale R, Tan YL, Iordanidou P, Pavón Arocas O, Claudi F, Stempel AV, Keshavarzi S, Petersen RS, Margrie TW, Branco T. A cortico-collicular circuit for orienting to shelter during escape. Nature. 2023 Jan;613(7942):111-119. doi: 10.1038/s41586-022-05553-9.

23. Caulfield MP. Muscarinic receptors--characterization, coupling and function. Pharmacol Ther. 1993;58(3):319-379. doi:10.1016/0163-7258(93)90027-b

24. Caulfield MP, Birdsall NJ. International Union of Pharmacology. XVII. Classification of muscarinic acetylcholine receptors. Pharmacol Rev. 1998;50(2):279–290.

25. Chen N, Sugihara H, Sur M. An acetylcholine-activated microcircuit drives temporal dynamics of cortical activity. Nat Neurosci. 2015 Jun;18(6):892–902. doi: 10.1038/nn.4002. Epub 2015 Apr 27. PMID: 25915477; PMCID: PMC4446146.

26. Chrastil ER, Sherrill KR, Aselcioglu I, Hasselmo ME, Stern CE. 2017. Individual Differences in Human Path Integration Abilities Correlate with Gray Matter Volume in Retrosplenial Cortex, Hippocampus, and Medial Prefrontal Cortex. eneuro. doi:10.1523/eneuro.0346-16.2017

27. Chrastil ER, Sherrill KR, Hasselmo ME, Stern CE. 2015. There and back again: Hippocampus and retrosplenial cortex track homing distance during human path integration. J Neurosci 35:15442–15452. doi:10.1523/JNEUROSCI.1209-15.2015

28. Clark BJ, Bassett JP, Wang SS, Taube JS. Impaired head direction cell representation in the anterodorsal thalamus after lesions of the retrosplenial cortex. J Neurosci. 2010;30(15):5289–5302. doi:10.1523/JNEUROSCI.3380-09.2010

29. Conner JM, Culberson A, Packowski C, Chiba AA, Tuszynski MH. Lesions of the Basal forebrain cholinergic system impair task acquisition and abolish cortical plasticity associated with motor skill learning. Neuron. 2003 Jun 5;38(5):819–29. doi: 10.1016/s0896-6273(03)00288-5. PMID: 12797965.

30. Conner JM, Kulczycki M, Tuszynski MH. Unique contributions of distinct cholinergic projections to motor cortical plasticity and learning. Cereb Cortex. 2010 Nov;20(11):2739–48. doi: 10.1093/cercor/bhq022. Epub 2010 Feb 24. PMID: 20181623; PMCID: PMC2951849.

31. Cooper BG, Mizumori SJY. 1999. Retrosplenial cortex inactivation selectively impairs navigation in darkness. Neuroreport 10:625–630.

32. Cho J, Sharp PE. Head direction, place, and movement correlates for cells in the rat retrosplenial cortex. Behav Neurosci. 2001;115(1):3–25. doi:10.1037/0735-7044.115.1.3

33. Cullen KE, Taube JS. Our sense of direction: progress, controversies and challenges. Nat Neurosci. 2017 Oct 26;20(11):1465–1473. doi: 10.1038/nn.4658. PMID: 29073639; PMCID: PMC10278035.

34. Dasari S, Hill C, Gulledge AT. A unifying hypothesis for M1 muscarinic receptor signalling in pyramidal neurons 2017. J Physiol. ;595(5):1711–1723. doi: 10.1113/JP273627.

35. Dembrow NC, Chitwood RA, Johnston D. 2010. Projection-specific neuromodulation of medial prefrontal cortex neurons. J Neurosci 30:16922–16937. doi:10.1523/JNEUROSCI.3644-10.2010

36. Derdikman D. Are the boundary-related cells in the subiculum boundary-vector cells? J Neurosci. 2009 Oct 28;29(43):13429–31. doi: 10.1523/JNEUROSCI.4176-09.2009.

37. Donoghue JP, Carroll KL. 1987. Cholinergic modulation of sensory responses in rat primary somatic sensory cortex. Brain Res 408:367–371. doi:10.1016/0006-8993(87)90407-0

38. Eckenstein FP, Baughman RW, Quinn J. 1988. An anatomical study of cholinergic innervation in rat cerebral cortex. Neuroscience 25:457–474. doi:10.1016/0306-4522(88)90251-5

39. Egan TM, North RA. Acetylcholine hyperpolarizes central neurones by acting on an M2 muscarinic receptor. Nature. 1986;319(6052):405-407. doi:10.1038/319405a0

40. Egorov A V., Hamam BN, Fransén E, Hasselmo ME, Alonso AA. 2002. Graded persistent activity in entorhinal cortex neurons. Nature 420:173–178. doi:10.1038/nature01171

41. Elduayen C, Save E. 2014. The retrosplenial cortex is necessary for path integration in the dark. Behav Brain Res 272:303–307. doi 10.1016/j.bbr.2014.07.009

42. Ferguson KA, Cardin JA. 2020. Mechanisms underlying gain modulation in the cortex. Nat Rev Neurosci. doi:10.1038/s41583-019-0253-y

43. Ferreira-Vieira TH, Guimaraes IM, Silva FR, Ribeiro FM. Alzheimer’s disease: Targeting the Cholinergic System. Curr Neuropharmacol. 2016;14(1):101–115. doi:10.2174/1570159x13666150716165726

44. Fu X, Ye H, Jia H, Wang X, Chomiak T, Luo F. 2019. Muscarinic acetylcholine receptor-dependent persistent activity of layer 5 intrinsic-bursting and regular-spiking neurons in primary auditory cortex. J Neurophysiol 122:2344–2353. doi:10.1152/jn.00184.2019

45. Ghosh M, Yang FC, Rice SP, Hetrick V, Gonzalez AL, Siu D, Brennan EKW, John TT, Ahrens AM, Ahmed OJ. Running speed and REM sleep control two distinct modes of rapid interhemispheric communication. Cell Rep. 2022 Jul 5;40(1):111028. doi: 10.1016/j.celrep.2022.111028. PMID: 35793619; PMCID: PMC9291430.

46. Gil Z, Connors BW, Amitai Y. Differential regulation of neocortical synapses by neuromodulators and activity. Neuron. 1997 Sep;19(3):679–86. doi: 10.1016/s0896-6273(00)80380-3.

47. Giovannini MG, Rakovska A, Benton RS, Pazzagli M, Bianchi L, Pepeu G. 2001. Effects of novelty and habituation on acetylcholine, GABA, and glutamate release from the frontal cortex and hippocampus of freely moving rats. Neuroscience 106:43–53. doi:10.1016/S0306-4522(01)00266-4

48. Goff KM, Liebergall SR, Jiang E, Somarowthu A, Goldberg EM. VIP interneuron impairment promotes in vivo circuit dysfunction and autism-related behaviors in Dravet syndrome [published online ahead of print, 2023 Jun 12]. Cell Rep. 2023;42(6):112628. doi:10.1016/j.celrep.2023.112628

49. Gulledge AT. 2024. Cholinergic Activation of Corticofugal Circuits in the Adult Mouse Prefrontal Cortex. J Neurosci. 44(**3**):e1388232023. doi: 10.1523/JNEUROSCI.1388-23.2023.

50. Gulledge AT, Bucci DJ, Zhang SS, Matsui M, Yeh HH. 2009. M1 receptors mediate cholinergic modulation of excitability in neocortical pyramidal neurons. J Neurosci 29:9888–9902. doi:10.1523/JNEUROSCI.1366-09.2009

51. Haj-Dahmane S, Andrade R. 1998. Ionic mechanism of the slow afterdepolarization induced by muscarinic receptor activation in rat prefrontal cortex. J Neurophysiol 80:1197–1210. doi:10.1152/jn.1998.80.3.1197

52. Hamlin AS, Windels F, Boskovic Z, Sah P, Coulson EJ. 2013. Lesions of the Basal Forebrain Cholinergic System in Mice Disrupt Idiothetic Navigation. PLoS One 8:53472. doi:10.1371/journal.pone.0053472

53. Hasselmo ME. Neuromodulation and cortical function: modeling the physiological basis of behavior. Behav Brain Res. 1995 Feb;67(1):1–27. doi: 10.1016/0166-4328(94)00113-t.

54. Hasselmo ME. Neuromodulation: acetylcholine and memory consolidation. Trends Cogn Sci. 1999 Sep;3(9):351–359. doi: 10.1016/s1364-6613(99)01365-0. PMID: 10461198.

55. Hasselmo ME. 2006. The role of acetylcholine in learning and memory. Curr Opin Neurobiol. doi:10.1016/j.conb.2006.09.002

56. Hasselmo ME. 2008. Grid cell mechanisms and function: Contributions of entorhinal persistent spiking and phase resetting. Hippocampus 18:1213–1229. doi:10.1002/hipo.20512

57. Hasselmo ME, Giocomo LM. Cholinergic modulation of cortical function. J Mol Neurosci. 2006;30(1-2):133–5. doi: 10.1385/JMN:30:1:133. PMID: 17192659.

58. Hattox AM, Nelson SB. 2007. Layer V neurons in mouse cortex projecting to different targets have distinct physiological properties. J Neurophysiol 98:3330–3340. doi:10.1152/jn.00397.2007

59. Hsieh CY, Cruikshank SJ, Metherate R. 2000. Differential modulation of auditory thalamocortical and intracortical synaptic transmission by cholinergic agonist. Brain Res 880:51–64. doi:10.1016/S0006-8993(00)02766-9

60. Hulse BK, Jayaraman V. Mechanisms Underlying the Neural Computation of Head Direction. Annu Rev Neurosci. 2020 Jul 8;43:31–54. doi: 10.1146/annurev-neuro-072116-031516. Epub 2019 Dec 24. PMID: 31874068.

61. Jacob PY, Casali G, Spieser L, Page H, Overington D, Jeffery K. An independent, landmark-dominated head-direction signal in dysgranular retrosplenial cortex. Nat Neurosci. 2017;20(2):173–175. doi:10.1038/nn.4465

62. Jiang L, Kundu S, Lederman JD, López-Hernández GY, Ballinger EC, Wang S, Talmage DA, Role LW. Cholinergic Signaling Controls Conditioned Fear Behaviors and Enhances Plasticity of Cortical-Amygdala Circuits. Neuron. 2016 Jun 1;90(5):1057–70. doi: 10.1016/j.neuron.2016.04.028. Epub 2016 May 5. PMID: 27161525; PMCID: PMC4891303.

63. Jiang X, Shen S, Cadwell CR, Berens P, Sinz F, Ecker AS, Patel S, Tolias AS. 2015. Principles of connectivity among morphologically defined cell types in adult neocortex. Science (80-) 350. doi:10.1126/science.aac9462

64. Jin J, Cheng J, Lee KW, Amreen B, McCabe KA, Pitcher C, Liebmann T, Greengard P, Flajolet M. 2019. Cholinergic neurons of the medial septum are crucial for sensorimotor gating. J Neurosci 39:5234–5242. doi:10.1523/JNEUROSCI.0950-18.2019

65. Jochems A, Reboreda A, Hasselmo ME, Yoshida M. 2013. Cholinergic receptor activation supports persistent firing in layer III neurons in the medial entorhinal cortex. Behav Brain Res 254:108–115. doi:10.1016/j.bbr.2013.06.027

66. Jochems A, Yoshida M. 2013. Persistent firing supported by an intrinsic cellular mechanism in hippocampal CA3 pyramidal cells. Eur J Neurosci 38:2250–2259. doi:10.1111/ejn.12236

67. Ju M, Gaussier P. A model of path integration and representation of spatial context in the retrosplenial cortex. Biol Cybern. 2020 Apr;114(2):303–313. doi: 10.1007/s00422-020-00833-x.

68. Kametani H, Kawamura H. Alterations in acetylcholine release in the rat hippocampus during sleep-wakefulness detected by intracerebral dialysis. Life Sci. 1990;47(5):421–426. doi:10.1016/0024-3205(90)90300-g

69. Kawaguchi Y. 1997. Selective cholinergic modulation of cortical GABAergic cell subtypes. J Neurophysiol 78:1743–1747. doi:10.1152/jn.1997.78.3.1743

70. Kawasaki H, Palmieri C, Avoli M. 1999. Muscarinic receptor activation induces depolarizing plateau potentials in bursting neurons of the rat subiculum. J Neurophysiol 82:2590–2601. doi:10.1152/jn.1999.82.5.2590

71. Keene CS, Bucci DJ. Contributions of the retrosplenial and posterior parietal cortices to cue-specific and contextual fear conditioning. Behav Neurosci. 2008 Feb;122(1):89–97. doi: 10.1037/0735-7044.122.1.89. PMID: 18298252.

72. Keshavarzi S, Bracey EF, Faville RA, et al. Multisensory coding of angular head velocity in the retrosplenial cortex. Neuron. 2022;110(3):532–543.e9. doi:10.1016/j.neuron.2021.10.031

73. Kilgard MP, Merzenich MM. 1998. Cortical map reorganization enabled by nucleus basalis activity. Science (80-) 279:1714–1718. doi:10.1126/science.279.5357.1714

74. Kimura F. 1999. Acetylcholine suppresses the spread of excitation in the visual cortex revealed by optical recording: Possible differential effect depending on the source of input. Eur J Neurosci 11:3597–3609. doi:10.1046/j.1460-9568.1999.00779.x

75. Kitanishi T, Umaba R, Mizuseki K. Robust information routing by dorsal subiculum neurons. Sci Adv. 2021 Mar 10;7(11):eabf1913. doi: 10.1126/sciadv.abf1913.

76. Knauer B, Jochems A, Valero-Aracama MJ, Yoshida M. 2013. Long-lasting intrinsic persistent firing in rat CA1 pyramidal cells: A possible mechanism for active maintenance of memory. Hippocampus 23:820–831. doi:10.1002/hipo.22136

77. Knierim JJ, Zhang K. Attractor dynamics of spatially correlated neural activity in the limbic system. Annu. Rev. Neurosci. 35, 267–285 (2012).

78. Kruglikov I, Rudy B. Perisomatic GABA release and thalamocortical integration onto neocortical excitatory cells are regulated by neuromodulators. Neuron. 2008 Jun 26;58(6):911–24. doi: 10.1016/j.neuron.2008.04.024. PMID: 18579081; PMCID: PMC2572574

79. Kurotani T, Miyashita T, Wintzer M, et al. Pyramidal neurons in the superficial layers of rat retrosplenial cortex exhibit a late-spiking firing property. Brain Struct Funct. 2013;218(1):239–254. doi:10.1007/s00429-012-0398-1

80. Lee SM, Seol JM, Lee I. Subicular neurons represent multiple variables of a hippocampal-dependent task by using theta rhythm. PLoS Biol. 2022;20(1):e3001546. Published 2022 Jan 31. doi:10.1371/journal.pbio.3001546

81. Lei, Y. T., Thuault, S. J., Launay, P., Margolskee, R. F., Kandel, E. R., & Siegelbaum, S. A. 2014. Differential contribution of TRPM4 and TRPM5 nonselective cation channels to the slow afterdepolarization in mouse prefrontal cortex neurons. Frontiers in cellular neuroscience, 8: 267. doi: 10.3389/fncel.2014.00267

82. Lein ES, Hawrylycz MJ, Ao N, et al. Genome-wide atlas of gene expression in the adult mouse brain. Nature. 2007;445(7124):168-176. doi:10.1038/nature05453

83. Lever C, Burton S, Jeewajee A, O’Keefe J, Burgess N. Boundary vector cells in the subiculum of the hippocampal formation. J Neurosci. 2009 Aug 5;29(31):9771–7. doi: 10.1523/JNEUROSCI.1319-09.2009. PMID: 19657030; PMCID: PMC2736390.

84. Levey AI. 1993. Immunological localization of m1-m5 muscarinic acetylcholine receptors in peripheral tissues and brain. Life Sci 52:441–448. doi:10.1016/0024-3205(93)90300-R

85. Leung LS, Péloquin P. Cholinergic modulation differs between basal and apical dendritic excitation of hippocampal CA1 pyramidal cells. Cereb Cortex. 2010;20(8):1865–1877. doi:10.1093/cercor/bhp251

86. Ljubojevic V, Luu P, Gill PR, Beckett LA, Takehara-Nishiuchi K, De Rosa E. Cholinergic Modulation of Frontoparietal Cortical Network Dynamics Supporting Supramodal Attention. J Neurosci. 2018 Apr 18;38(16):3988–4005. doi: 10.1523/JNEUROSCI.2350-17.2018. Epub 2018 Mar 23. PMID: 29572433; PMCID: PMC6705925

87. Lomi E, Mathiasen ML, Cheng HY, Zhang N, Aggleton JP, Mitchell AS, Jeffery KJ. Evidence for two distinct thalamocortical circuits in retrosplenial cortex. Neurobiol Learn Mem. 185:107525. doi: 10.1016/j.nlm.2021.107525.

88. Luchicchi A, Bloem B, Viaña JNM, Mansvelder HD, Role LW. 2014. Illuminating the role of cholinergic signaling in circuits of attention and emotionally salient behaviors. Front Synaptic Neurosci. doi:10.3389/fnsyn.2014.00024

89. Luntz-Leybman V, Bickford PC, Freedman R. 1992. Cholinergic gating of response to auditory stimuli in rat hippocampus. Brain Res 587:130–136. doi:10.1016/0006-8993(92)91437-J

90. Magistretti J, Ma L, Shalinsky MH, Lin W, Klink R, Alonso A. 2004. Spike Patterning by Ca2+-Dependent Regulation of a Muscarinic Cation Current in Entorhinal Cortex Layer II Neurons. J Neurophysiol 92:1644–1657. doi:10.1152/jn.00036.2004

91. Marrosu F, Portas C, Mascia MS, et al. Microdialysis measurement of cortical and hippocampal acetylcholine release during sleep-wake cycle in freely moving cats. Brain Res. 1995;671(2):329–332. doi:10.1016/0006-8993(94)01399-3

92. McKinney M, Coyle JT, Hedreen JC. 1983. Topographic analysis of the innervation of the rat neocortex and hippocampus by the basal forebrain cholinergic system. J Comp Neurol 217:103–121. doi:10.1002/cne.902170109

93. McNaughton BL, Battaglia FP, Jensen O, Moser EI, Moser MB. 2006. Path integration and the neural basis of the “cognitive map.” Nat Rev Neurosci. doi:10.1038/nrn1932

94. McNaughton BL, Chen LL, Markus EJ. 1991. “‘Dead reckoning,’” landmark learning, and the sense of direction: A neurophysiological and computational hypothesis. J Cogn Neurosci. doi:10.1162/jocn.1991.3.2.190

95. Mesulam MM, Mufson EJ, Wainer BH, Levey AI. 1983. Central cholinergic pathways in the rat: An overview based on an alternative nomenclature (Ch1-Ch6). Neuroscience 10:1185–1201. doi:10.1016/0306-4522(83)90108-2

96. Minces V, Pinto L, Dan Y, Chiba AA. 2017. Cholinergic shaping of neural correlations. Proc Natl Acad Sci U S A 114:5725–5730. doi:10.1073/pnas.1621493114

97. Mitsushima D, Sano A, Takahashi T. A cholinergic trigger drives learning-induced plasticity at hippocampal synapses. Nat Commun. 2013;4:2760. doi: 10.1038/ncomms3760. PMID: 24217681; PMCID: PMC3831287.

98. Mizuno H, Ikegaya Y. Late-spiking retrosplenial cortical neurons are not synchronized with neocortical slow waves in anesthetized mice. Neurosci Res. 2024 Jan 14:S0168-0102(24)00004-X. doi: 10.1016/j.neures.2024.01.001.

99. Muñoz W, Rudy B. Spatiotemporal specificity in cholinergic control of neocortical function. Curr Opin Neurobiol. 2014;26:149–160. doi:10.1016/j.conb.2014.02.015

100. Muñoz W, Tremblay R, Levenstein D, Rudy B. Layer-specific modulation of neocortical dendritic inhibition during active wakefulness. Science. 2017 Mar 3;355(6328):954–959. doi: 10.1126/science.aag2599. PMID: 28254942.

101. Nakajima Y, Nakajima S, Leonard RJ, Yamaguchi K. Acetylcholine raises excitability by inhibiting the fast transient potassium current in cultured hippocampal neurons. Proc Natl Acad Sci U S A. 1986 May;83(9):3022–6. doi: 10.1073/pnas.83.9.3022. PMID: 3010326; PMCID: PMC323439.

102. Navaroli VL, Zhao Y, Boguszewski P, Brown TH. 2012. Muscarinic receptor activation enables persistent firing in pyramidal neurons from superficial layers of dorsal perirhinal cortex. Hippocampus 22:1392–1404. doi:10.1002/hipo.20975

103. Nitzan N, McKenzie S, Beed P, English DF, Oldani S, Tukker JJ, Buzsáki G, Schmitz D. 2020. Propagation of hippocampal ripples to the neocortex by way of a subiculum-retrosplenial pathway. Nat Commun 11:1–17. doi:10.1038/s41467-020-15787-8

104. Olson JM, Tongprasearth K, Nitz DA. Subiculum neurons map the current axis of travel. Nat Neurosci. 2017 Feb;20(2):170–172. doi: 10.1038/nn.4464.

105. Opalka AN, Wang D V. 2020. Hippocampal efferents to retrosplenial cortex and lateral septum are required for memory acquisition. Learn Mem 27:310–318. doi:10.1101/LM.051797.120

106. Parikh V, Kozak R, Martinez V, Sarter M. 2007. Prefrontal Acetylcholine Release Controls Cue Detection on Multiple Timescales. Neuron 56:141–154. doi:10.1016/j.neuron.2007.08.025

107. Picciotto MR, Higley MJ, Mineur YS. 2012. Acetylcholine as a Neuromodulator: Cholinergic Signaling Shapes Nervous System Function and Behavior. Neuron. doi:10.1016/j.neuron.2012.08.036

108. Pinto L, Goard MJ, Estandian D, Xu M, Kwan AC, Lee SH, Harrison TC, Feng G, Dan Y. 2013. Fast modulation of visual perception by basal forebrain cholinergic neurons. Nat Neurosci 16:1857–1863. doi:10.1038/nn.3552

109. Polack PO, Friedman J, Golshani P. 2013. Cellular mechanisms of brain state-dependent gain modulation in visual cortex. Nat Neurosci 16:1331–1339. doi:10.1038/nn.3464

110. Rahman J, Berger T. 2011. Persistent activity in layer 5 pyramidal neurons following cholinergic activation of mouse primary cortices. Eur J Neurosci 34:22–30. doi:10.1111/j.1460-9568.2011.07736.x

111. Rasmusson DD. 2000. The role of acetylcholine in cortical synaptic plasticity. Behav Brain Res 115:205–218. doi:10.1016/S0166-4328(00)00259-X

112. Ratté S, Karnup S, Prescott SA. 2018. Nonlinear relationship between spike-dependent calcium influx and TRPC channel activation enables robust persistent spiking in neurons of the anterior cingulate cortex. J Neurosci 38:1788–1801. doi:10.1523/JNEUROSCI.0538-17.2018

113. Riekkinen P, Kuitunen J, Riekkinen M. 1995. Effects of scopolamine infusions into the anterior and posterior cingulate on passive avoidance and water maze navigation. Brain Res 685:46–54. doi:10.1016/0006-8993(95)00422-M

114. Roach JP, Eniwaye B, Booth V, Sander LM, Zochowski MR. Acetylcholine Mediates Dynamic Switching Between Information Coding Schemes in Neuronal Networks. Front Syst Neurosci. 2019;13:64. Published 2019 Nov 12. doi:10.3389/fnsys.2019.00064

115. Robles RM, Domínguez-Sala E, Martínez S, Geijo-Barrientos E. Layer 2/3 Pyramidal Neurons of the Mouse Granular Retrosplenial Cortex and Their Innervation by Cortico-Cortical Axons. Front Neural Circuits. 2020;14:576504. Published 2020 Nov 3. doi:10.3389/fncir.2020.576504

116. Sarter M, Hasselmo ME, Bruno JP, Givens B. 2005. Unraveling the attentional functions of cortical cholinergic inputs: Interactions between signal-driven and cognitive modulation of signal detection. Brain Res Rev. doi:10.1016/j.brainresrev.2004.08.006

117. Sarter M, Parikh V, Howe WM. Phasic acetylcholine release and the volume transmission hypothesis: time to move on. Nat Rev Neurosci. 2009 May;10(5):383–90. doi: 10.1038/nrn2635. PMID: 19377503; PMCID: PMC2699581.

118. Sarter M, Lustig C. 2020. Forebrain cholinergic signaling: Wired and phasic, not tonic, and causing behavior. J Neurosci. doi:10.1523/JNEUROSCI.1305-19.2019

119. Sarter M, Lustig C, Howe WM, Gritton H, Berry AS. 2014. Deterministic functions of cortical acetylcholine. Eur J Neurosci 39:1912–1920. doi:10.1111/ejn.12515

120. Savage LM. Sustaining high acetylcholine levels in the frontal cortex, but not retrosplenial cortex, recovers spatial memory performance in a rodent model of diencephalic amnesia. Behav Neurosci. 2012;126(2):226–236. doi:10.1037/a0027257

121. Sharp PE, Tinkelman A, Cho J. Angular velocity and head direction signals recorded from the dorsal tegmental nucleus of Gudden in the rat: implications for path integration in the head direction cell circuit. Behav. Neurosci. 115, 571–588 (2001).

122. Sherrill KR, Erdem UM, Ross RS, Brown TI, Hasselmo ME, Stern CE. 2013. Hippocampus and retrosplenial cortex combine path integration signals for successful navigation. J Neurosci 33:19304–19313. doi:10.1523/JNEUROSCI.1825-13.2013

123. Simonnet J, Brecht M. Burst Firing and Spatial Coding in Subicular Principal Cells. J Neurosci. 2019 May 8;39(19):3651–3662. doi: 10.1523/JNEUROSCI.1656-18.2019.

124. Solari N, Hangya B. 2018. Cholinergic modulation of spatial learning, memory and navigation. Eur J Neurosci. doi:10.1111/ejn.14089

125. Soma S, Shimegi S, Osaki H, Sato H. 2012. Cholinergic modulation of response gain in the primary visual cortex of the macaque. J Neurophysiol 107:283–291. doi:10.1152/jn.00330.2011

126. Soma S, Shimegi S, Suematsu N, Sato H. 2013. Cholinergic modulation of response gain in the rat primary visual cortex. Sci Rep 3:1–7. doi:10.1038/srep01138

127. Stackman RW, Taube JS. Firing properties of head direction cells in the rat anterior thalamic nucleus: dependence on vestibular input. J. Neurosci. 17, 4349–4358 1997.

128. Stancampiano R, Cocco S, Cugusi C, Sarais L, Fadda F. 1999. Serotonin and acetylcholine release response in the rat hippocampus during a spatial memory task. Neuroscience 89:1135–1143. doi:10.1016/S0306-4522(98)00397-2

129. Sullivan KE, Kraus L, Kapustina M, et al. Sharp cell-type-identity changes differentiate the retrosplenial cortex from the neocortex. Cell Rep. 2023;42(3):112206. doi:10.1016/j.celrep.2023.112206

130. Szymusiak R, Alam N, McGinty D. 2000. Discharge patterns of neurons in cholinergic regions of the basal forebrain during waking and sleep. Behav Brain Res 115:171–182. doi:10.1016/S0166-4328(00)00257-6

131. Tahvildari B, Fransén E, Alonso AA, Hasselmo ME. 2007. Switching between “on” and “off” states of persistent activity in lateral entorhinal layer III neurons. Hippocampus 17:257–263. doi:10.1002/hipo.20270

132. Taube JS. Head direction cells recorded in the anterior thalamic nuclei of freely moving rats. J. Neurosci. 15, 70–86 1995

133. Taube JS, Muller RU, Ranck JB Jr. Head-direction cells recorded from the postsubiculum in freely moving rats. I. Description and quantitative analysis. J Neurosci. 1990;10(2):420–435. doi:10.1523/JNEUROSCI.10-02-00420.1990

134. Taube JS, Muller RU. Comparisons of head direction cell activity in the postsubiculum and anterior thalamus of freely moving rats. Hippocampus. 1998;8(2):87–108. doi:10.1002/(SICI)1098-1063(1998)8:2<87::AID-HIPO1>3.0.CO;2-4

135. Tengelsen LA, Robertson RT, Yu J. 1992. Basal forebrain and anterior thalamic contributions to acetylcholinesterase activity in granular retrosplenial cortex of rats. Brain Res 594:10–18. doi:10.1016/0006-8993(92)91024-9

136. Thuault SJ, Malleret G, Constantinople CM, Nicholls R, Chen I, Zhu J, Panteleyev A, Vronskaya S, Nolan MF, Bruno R, Siegelbaum SA, Kandel ER. 2013. Prefrontal cortex HCN1 channels enable intrinsic persistent neural firing and executive memory function. J Neurosci 33:13583–13599. doi:10.1523/JNEUROSCI.2427-12.2013

137. Tsiola A, Hamzei-Sichani F, Peterlin Z, Yuste R. 2003. Quantitative morphologic classification of layer 5 neurons from mouse primary visual cortex. J Comp Neurol 461:415–428. doi:10.1002/cne.10628

138. Turner-Evans D, Wegener S, Rouault H, Franconville R, Wolff T, Seelig JD, Druckmann S, Jayaraman V. Angular velocity integration in a fly heading circuit. Elife. 2017 May 22;6:e23496. doi: 10.7554/eLife.23496. PMID: 28530551; PMCID: PMC5440168.

139. van der Goes MH, Voigts J, Newman JP, Toloza EHS, Brown NJ, Murugan P, Harnett MT. Coordinated head direction representations in mouse anterodorsal thalamic nucleus and retrosplenial cortex. Elife. 2024 Mar 12;13:e82952. doi: 10.7554/eLife.82952. PMID: 38470232; PMCID: PMC10932540.

140. van Groen T, Wyss JM. 2003. Connections of the retrosplenial granular b cortex in the rat. J Comp Neurol 463:249–263. doi:10.1002/cne.10757

141. van Groen T, Wyss JM. 1992. Connections of the retrosplenial dysgranular cortex in the rat. J Comp Neurol 315:200–216.

142. van Groen T, Wyss JM. 1990. Connections of the retrosplenial granular a cortex in the rat. J Comp Neurol 300:593–606. doi:10.1002/cne.903000412

143. Vann SD. A role for the head-direction system in geometric learning. Behav Brain Res. 2011;224(1):201–206. doi:10.1016/j.bbr.2011.05.033

144. Verhoog MB, Obermayer J, Kortleven CA, Wilbers R, Wester J, Baayen JC, De Kock CPJ, Meredith RM, Mansvelder HD. 2016. Layer-specific cholinergic control of human and mouse cortical synaptic plasticity. Nat Commun 7:1–13. doi:10.1038/ncomms12826

145. Vogt BA. 1984. Afferent specific localization of muscarinic acetylcholine receptors in cingulate cortex. J Neurosci 4:2191–2199. doi:10.1523/jneurosci.04-09-02191.1984

146. Wess J, Lambrecht G, Mutschler E, Brann MR, Dörje F. Selectivity profile of the novel muscarinic antagonist UH-AH 37 determined by the use of cloned receptors and isolated tissue preparations. Br J Pharmacol. 1991;102(1):246–250. doi:10.1111/j.1476-5381.1991.tb12161.x

147. Whishaw IQ. 1985. Cholinergic receptor blockade in the rat impairs locale but not taxon strategies for place navigation in a swimming pool. Behav Neurosci 99:979–1005. doi:10.1037/0735-7044.99.5.979

148. Winkler J, Suhr ST, Gage FH, Thal LJ, Fisher LJ. 1995. Essential role of neocortical acetylcholine in spatial memory. Nature 375:484–487. doi:10.1038/375484a0

149. Wyss JM, van Groen T. 1992. Connections between the retrosplenial cortex and the hippocampal formation in the rat: A review. Hippocampus 2:1–11. doi:10.1002/hipo.450020102

150. Wyss JM, van Groen T, Sripanidkulchai K. 1990. Dendritic bundling in layer I of granular retrosplenial cortex: Intracellular labeling and selectivity of innervation. J Comp Neurol 295:33–42. doi:10.1002/cne.902950104

151. Yamada-Hanff J, Bean BP. 2013. Persistent sodium current drives conditional pacemaking in CA1 pyramidal neurons under muscarinic stimulation. J Neurosci 33:15011–15021. doi:10.1523/JNEUROSCI.0577-13.2013

152. Yamawaki N, Corcoran KA, Guedea AL, Shepherd GMG, Radulovic J. 2019a. Differential Contributions of Glutamatergic Hippocampal→Retrosplenial Cortical Projections to the Formation and Persistence of Context Memories. Cereb Cortex 29:2728–2736. doi:10.1093/cercor/bhy142

153. Yamawaki N, Li X, Lambot L, Ren LY, Radulovic J, Shepherd GM. 2019b. Long-range inhibitory intersection of a retrosplenial thalamocortical circuit by apical tuft-targeting CA1 neurons. Nat Neurosci 22:618–626. doi:10.1101/427179

154. Yan HD, Villalobos C, Andrade R. 2009. TRPC channels mediate a muscarinic receptor-induced afterdepolarization in cerebral cortex. J Neurosci 29:10038– 10046. doi:10.1523/JNEUROSCI.1042-09.2009

155. Yao Z, van Velthoven CTJ, Nguyen TN, et al. 2021. A taxonomy of transcriptomic cell types across the isocortex and hippocampal formation. Cell. 184(12):3222–3241.e26. doi:10.1016/j.cell.2021.04.021

156. Yoder RM, Chan JHM, Taube JS. Acetylcholine contributes to the integration of self-movement cues in head direction cells. Behav Neurosci. 2017 Aug;131(4):312–24. doi: 10.1037/bne0000205.

157. Yoshida M, Fransén E, Hasselmo ME. 2008. mGluR-dependent persistent firing in entorhinal cortex layer III neurons. Eur J Neurosci 28:1116–1126. doi:10.1111/j.1460-9568.2008.06409.x

158. Yoshida M, Hasselmo ME. 2009. Persistent firing supported by an intrinsic cellular mechanism in a component of the head direction system. J Neurosci 29:4945– 4952. doi:10.1523/JNEUROSCI.5154-08.2009

159. Yoshida M, Knauer B, Jochems A. 2012. Cholinergic modulation of the CAN current may adjust neural dynamics for active memory maintenance, spatial navigation and time-compressed replay. Front Neural Circuits 6:10. doi:10.3389/fncir.2012.00010

160. Yousuf H, Nye AN, Moyer, Jr. JR. 2020. Heterogeneity of Neuronal Firing Type and Morphology in Retrosplenial Cortex of Male F344 Rats. J Neurophysiol 123. doi:10.1152/jn.00577.2019

161. Yuan R, Biswal BB, Zaborszky L. Functional Subdivisions of Magnocellular Cell Groups in Human Basal Forebrain: Test-Retest Resting-State Study at Ultra-high Field, and Meta-analysis. Cereb Cortex. 2019 Jul 5;29(7):2844–2858. doi: 10.1093/cercor/bhy150

162. Zannone S, Brzosko Z, Paulsen O, Clopath C. 2018. Acetylcholine-modulated plasticity in reward-driven navigation: a computational study. Sci Rep 8:1–20. doi:10.1038/s41598-018-27393-2

163. Zhang Z, Seguela P. 2010. Metabotropic Induction of Persistent Activity in Layers II/III of Anterior Cingulate Cortex. Cereb Cortex 20:2948–2957. doi:10.1093/cercor/bhq043

